# Resting-state theta oscillations and reward sensitivity in risk taking

**DOI:** 10.1101/2021.02.23.432425

**Authors:** Maria Azanova, Maria Herrojo Ruiz, Alexis V. Belianin, Vasily Klucharev, Vadim V. Nikulin

## Abstract

Females demonstrate greater risk aversion than males on a variety of tasks, but the underlying neurobiological basis is still unclear. We studied how theta (4-7 Hz) oscillations at rest related to three different measures of risk taking. Thirty-five participants (15 females) completed the Bomb Risk Elicitation Task (BRET), which allowed us to measure risk taking during an economic game. The Domain-Specific Risk-Taking Scale (DOSPERT) was used to measure self-assessed risk attitudes as well as reward and punishment sensitivities. In addition, the Barratt Impulsiveness Scale (BIS11) was included to quantify impulsiveness. To obtain measures of frontal theta asymmetry and frontal theta power, we used magnetoencephalography (MEG) acquired prior to task completion, while participants were at rest. Frontal theta asymmetry correlated with average risk taking during the game but only in the female sample. By contrast, frontal theta power correlated with risk taking as well as with measures of reward and punishment sensitivity in the joint sample. Importantly, we showed that reward sensitivity mediated a correlation between risk taking and the power of theta oscillations localized to the anterior cingulate cortex. In addition, we observed significant sex differences in source- and sensor-space theta power, risk taking during the game, and reward sensitivity. Our findings suggest that sensitivity to rewards, associated with resting-state theta oscillations in the anterior cingulate cortex, is a trait that potentially contributes to sex differences in risk taking.

## 1 Introduction

Behavioral heterogeneity is a pervasive feature of risk taking and decision-making. A neural trait approach suggests that heterogeneity in behavior can be at least partially explained by stable brain-based characteristics of individuals (Nash & Knoch, 2016). It was reported that, on average, females take fewer risks than males (e.g., Jianakoplos & Bernasek, 1998; Charness & Gneezy, 2012). This study investigated neural traits in relation to an interindividual variability in risk preferences in a sample containing both males and females. The consistent sex-related difference in risk preferences suggests the existence of sex-specific neural traits associated with risk attitudes (Ball, Balthazart, & McCarthy, 2014). Therefore, we examined if variability in risk attitudes among participants of both sexes could be explained by considering only brain-based measures and without accounting for their sex per se.

On the neural level, electroencephalography (EEG) studies found positive correlations between resting-state right-left frontal theta (4-7 Hz) asymmetry (rsFTA) and risk taking during an economic task (Gianotti et al., 2009; Studer, Pedroni, & Rieskamp, 2013). An association between the activity of the frontal lobes and trait behavioral inhibition may explain these results (e.g., Garavan, Ross, & Stein, 1999; Aron, Robbins, & Poldrack, 2004; Schiller, Gianotti, Nash, & Knoch, 2014). In particular, previous studies suggested that the level of risk aversion may reflect how well one suppresses an urge to go for a riskier, more tempting option. Noticeably, these studies examined neural signatures of risk taking in exclusively (Gianotti et al., 2009) or mostly (70%; Studer et al., 2013) female samples. However, there is recent evidence that males and females do not significantly differ in frontal EEG asymmetries across various frequency bands, including the theta band (Ocklenburg et al., 2019). Therefore, it remains unclear whether the observed sex differences in risk attitudes are related to rsFTA at all and whether rsFTA correlates with risk taking in a joint sample (i.e., a sample containing both males and females as opposed to participants of one sex).

Accordingly, our first goal was to determine whether rsFTA correlated with risk preferences in the joint sample. In this context, we aimed to replicate previous EEG findings for female and joint samples (Gianotti et al., 2009; Studer et al., 2013) but using magnetoencephalography (MEG) recordings and a new paradigm from behavioral economics. Following previous studies (Kamarajan et al., 2008; Gianotti et al., 2009; Gianotti et al., 2012; Lee & Jeong, 2013), we also examined if neuronal activity and risk taking were associated with self-assessed measures of impulsivity.

Our second goal was to determine whether resting-state frontal theta power (rsFT) could be an alternative neural trait underlying risk attitudes. Massar, Rossi, Schutter, & Kenemans (2012) demonstrated that resting-state theta/beta ratio correlated with feedback-related negativity (FRN) and subsequent disadvantageous/risky behavior during a gambling task in a sample that included both males and females. However, this result was only significant in a subsample with high punishment sensitivity scores. The follow-up study (Massar, Kenemans, & Schutter, 2014) further found that resting-state theta oscillations predicted reinforcement learning during the Iowa Gambling Task (IGT, Bechara, Damasio, Damasio, & Anderson, 1994) and correlated with reward sensitivity in the joint sample. In particular, Massar et al. (2014) showed that higher theta power at frontal and central sites was associated with choices from high-reward/high-loss (disadvantageous) decks. Furthermore, reactions to losses and gains have previously been linked to in-task changes in theta oscillations (e.g., Cohen, Elger, & Ranganath, 2007; Kamarajan et al., 2008; Cavanagh, Frank, Klein, & Allen, 2010; Crowley et al., 2014). Based on these findings, we hypothesized that rsFT would be positively correlated with risk taking and with self-assessed measures of reward sensitivity in the joint sample.

Finally, we aimed to analyze whether resting-state theta oscillations localized to the anterior cingulate cortex (ACC) correlated with risk attitudes. ACC was chosen as the region of interest for three reasons. First, various functional magnetic resonance imaging (fMRI) studies revealed its involvement in risk taking (e.g., Christopoulos, Tobler, Bossaerts, Dolan, & Schultz, 2009; Engelmann & Tamir, 2009; Fukunaga, Purcell, & Brown, 2018). Second, activity in ACC has been associated with frontal theta oscillations (Scheeringa et al., 2008; Massar et al., 2012). Third, this region may be related to sex differences in decision-making. Santesso, Dzyundzyak, & Segalowitz (2011) found sex differences in the FRN and reward and punishment sensitivities - all these measures also correlated with ACC activity. An fMRI study by Zhou et al. (2014) demonstrated that males and females differed in the baseline brain activity associated with risk attitudes. In particular, sex differences were found in regions of the default mode network, including ACC. Accordingly, ACC-related theta oscillations were a strong candidate to explain sex differences in risk taking.

To measure risk taking, we used the Bomb Risk Elicitation Task (BRET, Crosetto & Filippin, 2013). In one trial of this task, participants decide how many boxes to collect out of 100. Each of these boxes has the same probability of containing a bomb. The gain increases linearly with the number of boxes collected, but a participant wins nothing if the bomb is among the collected boxes. Thus, the task is framed entirely in the gain domain. Because probabilities of winning and possible outcomes of each choice are accessible to participants, the BRET measures specifically attitudes towards risk as opposed to ambiguity – the kind of uncertainty when probability distribution of possible outcomes is unknown (Huettel, Stowe, Gordon, Warner, & Platt, 2006). Consequently, there is no learning in this task, because it has a static structure that is explained to participants from the beginning, and, therefore, single-trial changes in risk preferences reflect state-like behavioral variability, unlike in the IGT. Importantly, BRET requires minimal numeracy skills and, from a theoretical-economic perspective, is not affected by the degree of loss aversion (increased weighting of possible losses as opposed to possible gains, Kahneman & Tversky, 1979), which could otherwise bias estimates of risk attitudes. The task also avoids discontinuity in risk-attitude measurement because it has finer dimensionality (101 choices in one trial) as compared to the Devil’s Task (7 choices in one trial) used previously by Gianotti et al. (2009). Moreover, in BRET, as opposed to both the Devil’s Task (Slovic, 1966) and the Balloon Analogue Risk Task (BART, Lejuez et al., 2002), a trial is not interrupted when a participant makes a no-win choice (finds a bomb): they finish the selection, revealing their true preference, and only then feedback is provided. This enables avoiding the truncation of data, especially for estimates of high-risk choices. Notably, Pedroni et al., 2017 observed that distinct measures of risk taking are associated with different ‘cognitive strategies’. We thus aimed to address this aspect by examining three distinct measures of risk taking. Apart from measures based on game performance, we used the Domain-Specific Risk-Taking Scale (DOSPERT, Blais & Weber, 2006), which measures self-assessed likelihood to take risks as well as punishment and reward sensitivities to risky actions across several decision-making domains.

## 2 Materials and methods

### 2.1 Participants

We recruited 35 right-handed individuals (15 females; average age females = 21.93, SD = 2.96; average age males = 22.55, SD = 3.95; no significant age difference) without a history of psychiatric or neurological disorders and any metal in the body. All participants had normal or corrected to normal vision. All of them were native Russian speakers. According to the power analysis, to reliably observe a correlation of 0.45 (comparable to results of the previous study by Gianotti et al., 2009) with power 0.8 and confidence 0.95, 35-36 observations were needed.

Participants were recruited via social media. The experiment was carried out in accordance with the recommendations of the Declaration of Helsinki and its amendments, and the protocol was approved by the ethics committee of the National Research University Higher School of Economics. Data collection took place at the Center for Neurocognitive Research, Moscow State University of Psychology and Education (MEG Center). A signed consent form was obtained from all participants at the beginning of the experimental session.

### 2.2 Behavioral Procedures

After the instructions, participants went through two blocks of 7-minute eyes-closed resting-state recordings with MEG. Here the participants were instructed to relax and to sit still.

Next, participants performed a modified version of the dynamic BRET (Crosetto & Filippin, 2013; Holzmeister, & Pfurtscheller, 2016). The game had 30 trials and it lasted around 10-15 minutes in total, depending on a participant’s speed and strategy. In each trial, a participant was presented with a 10-by-10 matrix that contained 100 boxes (Figure 1). She/he could select them sequentially one-by-one from the upper left corner to the bottom right corner. During the game, participants had to press the green button with the right-hand index finger to open a subsequent box and press the blue button with the right-hand middle finger to end a trial. Participants were not aware of the exact number of trials in the game. If one of the selected boxes contained a bomb, a participant won nothing in a trial. If none of the selected boxes contained a bomb, then a participant received 10 rubles for each chosen box. The bomb’s location was determined randomly in each trial and participants were informed about it during the instructions. Participants were notified whether there is a bomb among selected boxes after they chose to end a trial. This was done to avoid truncation of data. The outcome was presented on a separate screen after the participant had decided to stop the selection. The feedback screen revealed the number of selected boxes (‘You selected X boxes’), the location of the bomb (‘The bomb was in a cell X’), and the outcome (‘You won Z rubles’ or ‘You won nothing’).

**Figure 1.**
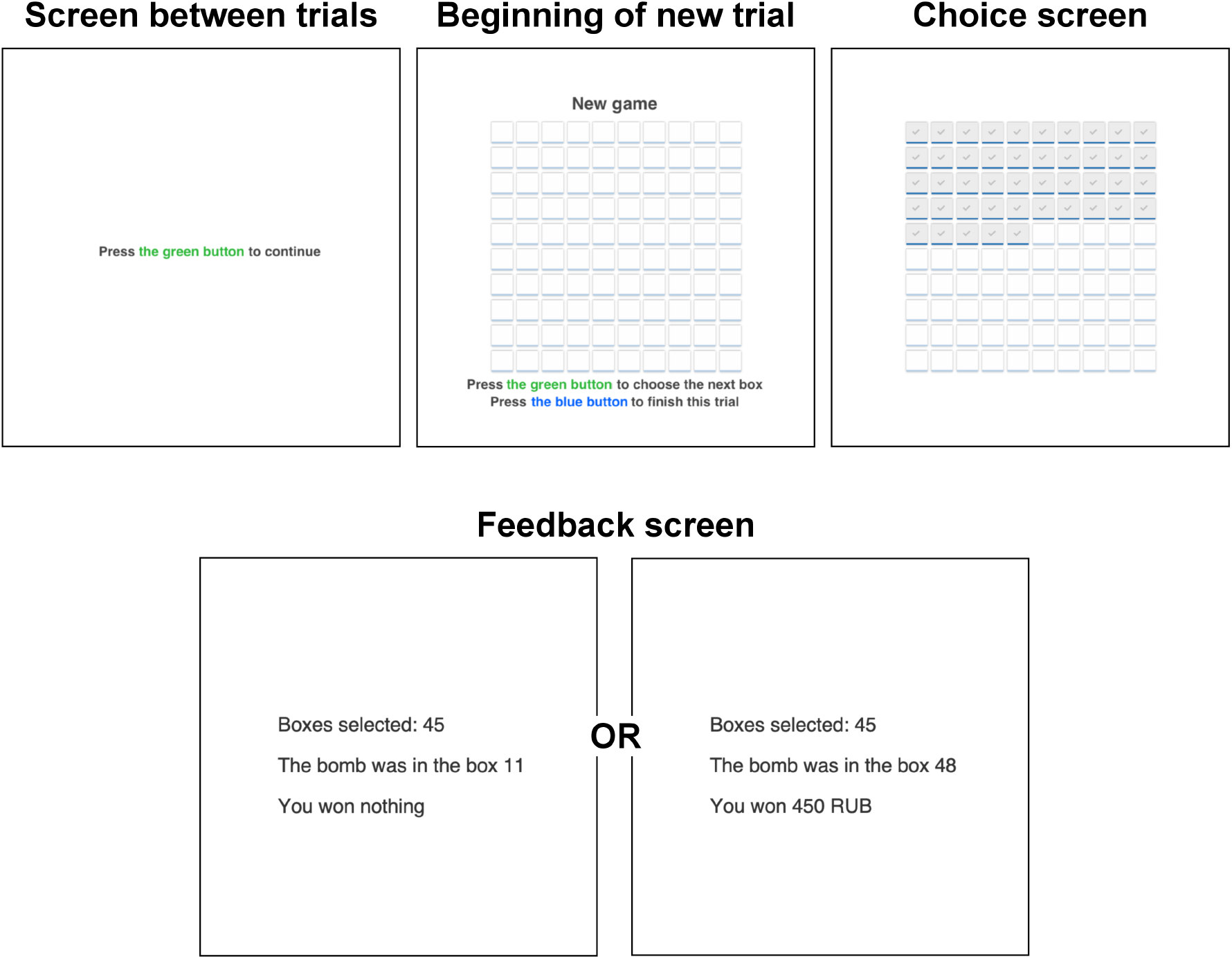
One trial of the BRET (the Bomb Risk Elicitation Task) adapted for the study.

After completing the task, participants went through another two 7-minute blocks of eyes-closed resting-state recordings with MEG. Overall, participants spent around 40 minutes in MEG’s shielded room.

After the MEG session, participants filled the following questionnaires in a separate room: the Domain-Specific Risk-Taking Scale (30-item version DOSPERT, Blais & Weber, 2006), and Barratt Impulsiveness Scale (BIS11, Patton, Stanford, & Barratt, 1995). The BRET and the questionnaires were programmed using PsychoPy software (Peirce, 2007). Each session lasted approximately 1-1.5 hours, including preparation and instructions (Figure 2).

**Figure 2.**
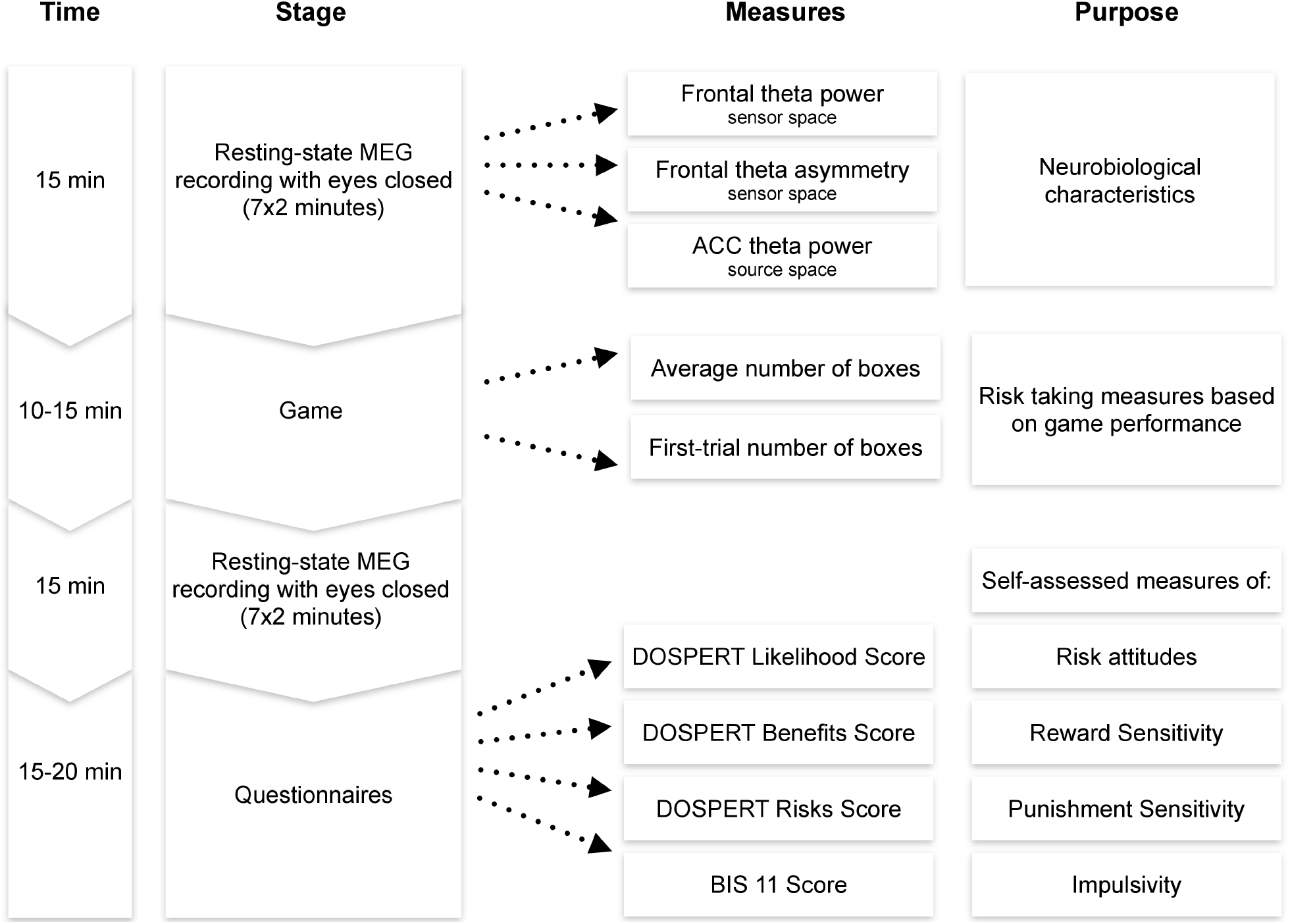
Experimental procedure. MEG - magnetoencephalography. ACC - anterior cingulate cortex. DOSPERT - Domain-Specific Risk-Taking Scale. BIS - Barratt Impulsiveness Scale.

Participants received 500 Russian rubles for participation and an additional bonus. The bonus was based on the outcome of one randomly selected trial, which was also made clear during the instructions. The bonus varied between 0 Russian rubles (18 participants) and 600 Russian rubles (17 participants; average = 325, SD= 106).

### 2.3 MEG Recording

The data was acquired with 306-channel magnetoencephalography ‘Neuromag VectorView’ (Elekta, Finland) consisting of 204 planar gradiometers and 102 magnetometers. The sampling frequency was 1000 Hz. The filter settings during the data recording were: lowpass 330 Hz, highpass 0.10 Hz. We controlled the head movement using a head-position indicator (HPI) with coils attached to the scalp.

In addition, two bipolar electrooculograms (EOG, vertical and horizontal) and a bipolar electrocardiogram (ECG) were recorded.

### 2.4 Behavioral Data Analysis

Two measures of risk attitude were computed based on the BRET performance: (1) the average number of boxes chosen in all trials of the game; (2) the number of boxes chosen in the first trial.

We studied the average behavior in the game because resting-state theta oscillations were previously associated with risk taking using average performance in tasks with repeated trials (Schutter & Van Honk 2005; Gianotti et al., 2009; Massar et al., 2012; Studer et al., 2013; Massar et al., 2014). We also studied behavior in the first trial of the game because it was previously suggested that there is no reference point in the first trial, while it could arise in subsequent trials based on previous performance (Crosetto & Filippin, 2013). The presence of a reference point potentially leads to an implicit loss aversion: a participant would consider zero win a loss if he/she expected to win a certain positive amount, which may bias risk attitude estimates (Koszegi & Rabin, 2006; Ert & Erev, 2013). Moreover, using several trials in the game may induce hedging or eventual boredom. Along with loss aversion, it may also lead to biases in estimates of risk attitudes based on average behavior, even if a participant receives payment only for one randomly chosen trial (Harrison, Martínez-Correa, & Swarthout, 2015; Cox, Sadiraj, & Schmidt, 2015). Nevertheless, Crosetto & Filippin (2016) stated that risk preferences in the repeated BRET were highly correlated with the one-shot version. It allowed us to assume that average behavior in the game would be a valid indicator of risk preferences even in the case of high intertrial variability of choices.

Crosetto & Filippin (2013) showed that the number of boxes chosen in the BRET was well suited for assessing risk preferences. A risk-neutral subject would choose 50 boxes out of 100 in each trial of a dynamic game because this strategy maximizes the objective expected winning amount. The fewer boxes are chosen - the more risk-averse one is. Risk-loving participants would ideally choose more than 50 boxes, and risk-averse participants would choose less than 50 boxes. For derivation of risk-attitude coefficients based on the number of boxes chosen in the BRET, please, refer to Crosetto & Filippin (2013). We discuss advantages of BRET as compared to other methods of assessing risk preferences in the Introduction.

Results of the Barratt Impulsiveness Scale (BIS11, Patton et al., 1995) were used as a self-assessed measure of impulsiveness following previous studies (e.g., Gianotti et al., 2009). The Domain-Specific Risk-Taking Scale (DOSPERT, Blais & Weber, 2006) has three subscales: it measures one’s propensity to participate in risky activities (‘how likely are you to..?’), as well as expected benefits (‘how beneficial is this?’) and perceived risks (‘how risky is this?’) of such activities. We used the DOSPERT likelihood subscale as an additional self-assessed measure of risk preferences, and the DOSPERT benefits and risks subscales as self-assessed measures of reward and punishment sensitivities to risky activities respectively. Even though DOSPERT includes 5 subscales relating to different domains of risk, such as financial or social, we did not consider them separately. Recent research shows that despite great inter- and intraindividual variability on these facets, there might be a more general underlying risk propensity that is also predictive of real-life behaviors (Highhouse, Nye, Zhang, & Rada, 2017). We did not include the BIS/BAS questionnaire (Carver & White, 1994) to quantify reward and punishment sensitivities. The rationale behind our decision was that these scores do not consistently differentiate motivational (reward and punishment sensitivity) and control (inhibition and impulsivity) components (Smillie, Jackson, & Dalgleish, 2006; Leone & Russo, 2009; Penolazzi, Gremigni, & Russo, 2012), and therefore do not straightforwardly relate to risk propensity across studies. By contrast, DOSPERT scores are more easily interpretable.

In addition, we used single-trial analysis to determine how participants changed their choices based on previous outcomes. We formalized this measure as a percent change in the number of boxes in a current trial as compared to a previous trial. Then, for each participant, we obtained two averaged measures of percent changes in the number of boxes after losing and winning. Even though the BRET does not presuppose learning, we considered these two additional measures as game-based indicators of punishment and reward sensitivities in the reinforcement learning sense. However, interpretations of outcome sensitivities based on DOSPERT subscales and these game-based measures differ significantly: DOSPERT scores quantify how pleasurable or undesirable participants find various risky activities, while percent change in the number of boxes in reaction to feedback indicates how risk preferences were affected by a previous outcome in the game. We provide results for game-based measures of reward and punishment sensitivities in Supplementary Table 1.

The behavioral data were processed using R software. In accordance with a previous protocol (Crosetto & Filippin, 2013), we excluded 9 trials from the analysis of a total of 1050 trials because 0 boxes (7 trials), 1 box (1 trial) or 2 boxes (1 trial) were selected in these trials. Among the remaining trials, the minimum value was 5 chosen boxes.

### 2.5 Sensor Space Analysis

The MEG data were pre-processed using the Elekta Neuromag software MaxFilter to compensate for head movement and interpolate bad channels, as well as to project noisy sources outside of the head. Next, we used the MNE-Python toolbox (Gramfort et al., 2014) to remove eye and heart-beat artifacts using bipolar EOG and ECG channels and independent component analysis (fastICA). In addition, we visually inspected the data for previously unaccounted artifacts (movement, system artifacts) to remove them before further processing.

Further analysis was carried out in MATLAB®. To study resting-state activity before the game, we chose the second 7-minutes eyes-closed resting-state recording. The first resting-state recording was excluded from analysis because it followed the start of the experiment. To ensure reliability of our results, we separately analyzed the MEG data from the resting-state recording after the game. For this, we chose the fourth 7-minutes resting-state recording, and we excluded the third recording because it followed the game and announcement of the final outcome immediately. One participant was excluded from this analysis due to technical problems with the MEG system. Therefore, the final sample for analysis of post-game resting-state activity included 34 participants (15 females). Detailed results of this separate analysis are provided in Supplementary Figures 2,3, and 5.

Consistently with our hypotheses outlined in the Introduction, the analysis of the MEG signals focused on the theta (4-7 Hz) power. We used magnetometer-based, rather than gradiometer-based, measurements in our analyses, because magnetometer data is more sensitive to deep sources such as those in ACC (Vrba & Robinson, 2001; Enatsu et al., 2008). The theta power was calculated as a mean of the squared signal obtained after bandpass filtering between 4 and 7 Hz (4th order, Butterworth filter). We then averaged the theta power across 26 sensors pertaining to regions of interest (ROIs) from the right and left frontal cortices (Figure 3).

**Figure 3.**
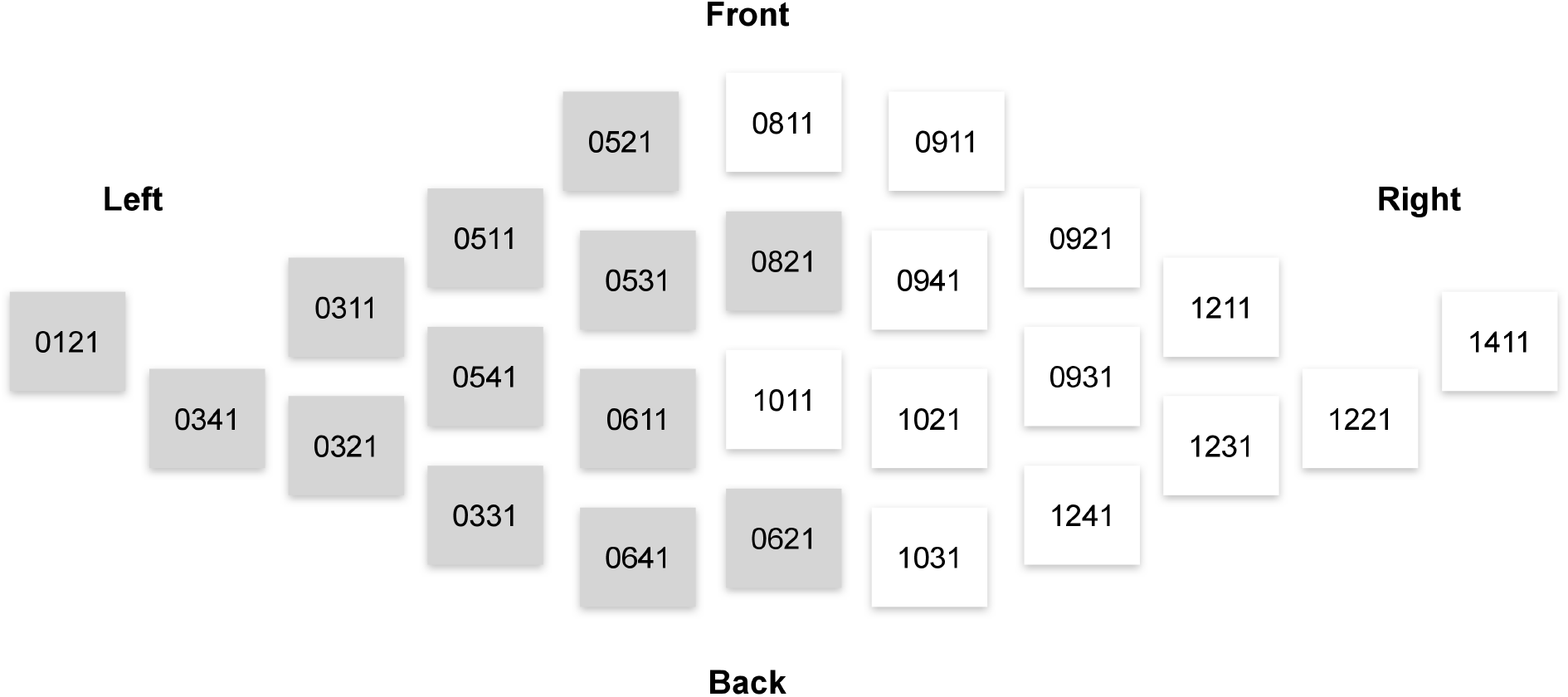
Elekta Neuromag sensors that were used in the analysis of sensor-space activity. Magnetometers used in computations of measures related to the left and right frontal lobes are denoted by grey and white, respectively.

To assess rsFTA, we followed a standard protocol used in previous EEG studies (Coan & Allen, 2004; Harmon-Jones & Gable, 2018; Ocklenburg et al., 2019). rsFTA is computed as the difference between the natural logarithm of theta power over the right and left frontal lobes: (ln[right]-ln[left]). This measure is on a scale where zero represents symmetrical activity, positive values represent greater right than left frontal activity, and negative values - greater relative left frontal activity. To obtain a measure representing rsFT, we summed theta power over the right and left frontal lobes: (right+left).

### 2.6 Source Space Analysis

Based on previous studies (Scheeringa et al., 2008; Massar et al., 2012), we expected that primarily theta power in the ACC would be associated with risk attitudes. Therefore, additional analyses in the source space were carried out to estimate theta power in the ACC using the Fieldtrip software (Oostenveld, Fries, Maris, & Schoffelen, 2011). For MRI segmentation, coregistration, and forward model estimation, we used a standard anatomical MRI template (“colin27”) from the FieldTrip toolbox. Forward modeling used a 5mm resolution grid, resulting in a source space of 38874 grid points (18693 inside the brain). Next, for inverse modeling, we reconstructed source space activity using Exact Low-Resolution Electromagnetic Tomography (eLORETA, Pascual-Marqui et al., 2011) based on magnetometer measurements (regularization parameter lambda=0.05). After inverse modeling, we extracted theta power averaged over time for each voxel. We then averaged the theta power across the voxels pertaining to the ACC ROI based on the MNI coordinates in the AAL-atlas (labels ‘Cingulum_Ant_L’ and ‘Cingulum_Ant_R’).

### 2.7 Statistical Analyses

We used R software to perform the statistical analyses. Measures related to frontal theta power and ACC theta power were standardized to undergo statistical analyses. Non-parametric Spearman correlations were used to examine the relationships between variables. To address the multiple comparisons problem, we implemented the Benjamini-Hochberg method to control the false discovery rate at level q = 0.05 (FDR; Benjamini & Hochberg, 1995). When reporting significant effects after controlling for the FDR, we provide the unadjusted p-value of the significant effect and denote that the result is significant after FDR control. Results that were not significant after controlling for FDR are followed by ‘n.s.’ To assess sex differences in behavioral or neural measures, we carried out two-sided independent 2-group Mann-Whitney U-tests. To confirm that specifically fronto-medial theta power is related to reward and punishment sensitivities (Cavanagh et al., 2010; Massaret al., 2014), we performed additional statistical analysis of the MEG data in sensor space using non-parametric cluster-based permutation tests on t-statistics (Maris & Oostenveld, 2007). Results of this analysis are in Supplementary Figures 4 and 5. Apart from performing non-parametric analysis, we provide results of regression analysis. It was included to test interaction effects and to compare how sex and neurobiological measures were related to measures of risk taking. Detailed regression results are in Supplementary Tables 2 and 3. To robustly determine if a given variable significantly improves performance of a linear model, we used ANOVA’s F-test to compare the two models: with and without a term of interest. Finally, we used packages mediation (Tingley, Yamamoto, Hirose, Keele, & Imai, 2014) and lavaan (Rosseel, 2012) to perform mediation analysis with one mediator and two mediators respectively. We tested the significance of indirect effects using bootstrapping procedures: unstandardized indirect effects were computed for each of 1000 bootstrapped samples.

## 3 Results

### 3.1 Behavioral Measures

#### 3.1.1 Risk Attitudes

On average, in all 30 trials, participants opened 44.86 boxes (SD = 9.92). According to the average behavior in the game, 2 participants were risk-neutral (number of boxes = 50), 20 participants were risk-averse (number of boxes < 50), 13 participants were risk-seeking (number of boxes > 50). In the first trial, participants opened, on average, 38.03 boxes (SD = 14.94). Average risk taking significantly correlated with risk taking in the first trial (Spearman’s ρ = 0.58, P = 0.0002, significant after FDR control) and with the self-assessed likelihood to take risks according to the DOSPERT likelihood subscale (Spearman’s ρ = 0.37, P = 0.03, FDR-controlled). However, first-trial risk taking and the DOSPERT likelihood subscale did not correlate significantly (Spearman’s ρ = 0.29, n.s.).

#### 3.1.2 Sex Differences in Risk Taking

On average, males (number of boxes = 48.38, SD = 8.66) took more risks than females (number of boxes = 40.18, SD = 9.79): Mann-Whitney U-test P = 0.01 (Figure 4). Males (number of boxes = 43.65, SD = 13.29) also chose significantly more boxes in the first trial of the game than females (number of boxes = 30.53, SD = 14.03): Mann-Whitney U-test P = 0.008. However, the two groups did not significantly differ according to the DOSPERT likelihood subscale: Mann-Whitney U-test P = 0.5.

**Figure 4.**
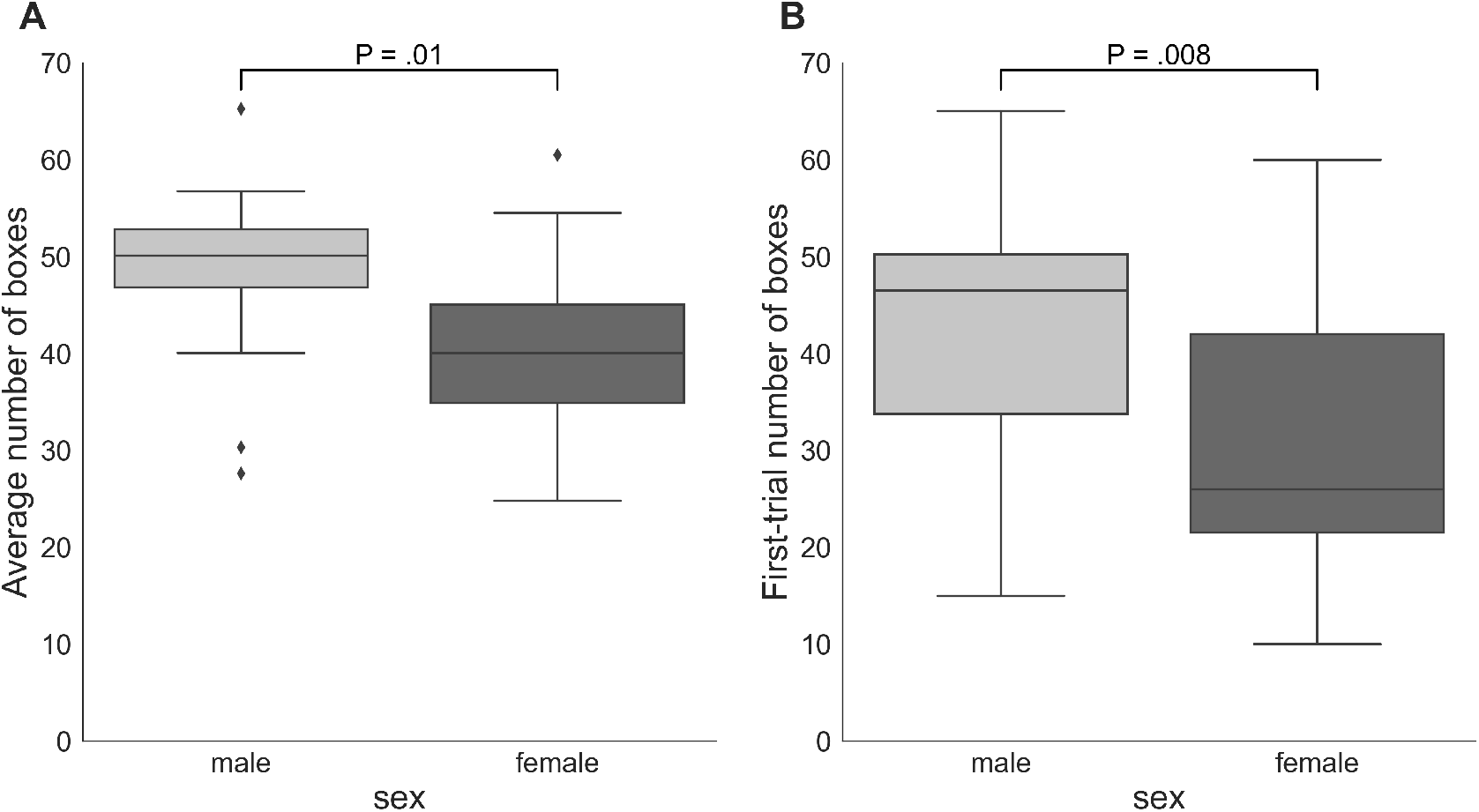
Box plots for the numbers of boxes chosen by male and female participants on average and during the first trial of the game. The proportion of the interquartile range (IQR) used to identify outliers equals 1.5. Outliers are denoted by diamond symbols. P - p-values of two-sided independent 2-group Mann-Whitney U-tests.

#### 3.1.3 Questionnaires

Among all DOSPERT and BIS11 subscales, only the DOSPERT benefits subscale positively correlated with the risk attitude measures obtained with BRET: with average (Spearman’s ρ = 0.47, P = 0.004, FDR-controlled) and first-trial (Spearman’s ρ = 0.55, P = 0.0006, FDR-controlled) risk taking in the game. There was a significant sex-related difference in the DOSPERT benefits scores (Mann-Whitney U-test P = 0.02). On average, males (score = 107.75, SD = 14.59) scored higher on self-assessed reward sensitivity than females (score = 96.33, SD = 15.27). DOSPERT subscales had high internal consistency as measured by Cronbach’s alpha (DOSPERT likelihood α = 0.84; DOSPERT benefits α = 0.72; DOSPERT risks α = 0.74), while BIS11 did not have high reliability (α = 0.43).

### 3.2 Frontal Theta Asymmetry (rsFTA)

Average risk taking during the BRET significantly correlated with rsFTA in the female subsample (Spearman’s ρ = 0.69, P = 0.004, FDR-controlled), but not in the male subsample (Spearman’s ρ = 0.28, n.s.) or the whole sample (Spearman’s ρ = 0.38, n.s.). Risk taking in the first trial of the game did not significantly correlate with rsFTA (Spearman’s ρ = 0.12, n.s.). There were no significant associations of rsFTA with first-trial risk taking in female (Spearman’s ρ = 0.25, n.s.) or male (Spearman’s ρ = −0.08, n.s.) subsamples. Correlations of rsFTA with DOSPERT likelihood scores were moderate, but not significant. Joint sample: Spearman’s ρ = 0.30, n.s. Female subsample: Spearman’s ρ = 0.35, n.s. Male subsample: Spearman’s ρ = 0.29, n.s. Females and males did not differ significantly in rsFTA (Mann-Whitney U-test P = 0.54). Finally, rsFTA did not significantly correlate with any of BIS11 subscales. The highest correlation of rsFTA was with the BIS11 self-control subscale (Spearman’s ρ = 0.4, n.s.). Thus, exclusively in the female subsample, we found a significant correlation between rsFTA and average risk taking during the game. This result was replicated based on the resting-state recording after the game: the correlation was significant only in the female subsample (Spearman’s ρ = 0.74, P = 0.001, FDR-controlled).

### 3.3 Frontal Theta Power (rsFT)

Correlation of average risk taking in the game with rsFT in the joint sample was not significant (Spearman’s ρ = 0.31, n.s.). By contrast, rsFT was significantly positively correlated with risk taking in the first trial of the game (Spearman’s ρ = 0.46, P = 0.01, FDR-controlled) as well as with the DOSPERT benefits subscale (Spearman’s ρ = 0.4, P = 0.02, FDR-controlled) and negatively correlated with the DOSPERT risks subscale (Spearman’s ρ = −0.4, P = 0.02, FDR-controlled) (Figure 5). We confirmed the significant results for DOSPERT subscales based on the resting-state recording after the game and also based on non-parametric statistical clustering in sensor space (see Supplementary Figures 2-5). The correlation of first-trial risk taking with rsFT after the game was not significant. However, we did observe significant clusters of midfrontal theta activity before and after the game that were associated with first-trial risk taking (Supplementary Figures 4 and 5). Furthermore, rsFT did not significantly correlate with DOSPERT likelihood scores. Last, males had significantly higher rsFT than females (Mann-Whitney U-test P = 0.0002).

**Figure 5.**
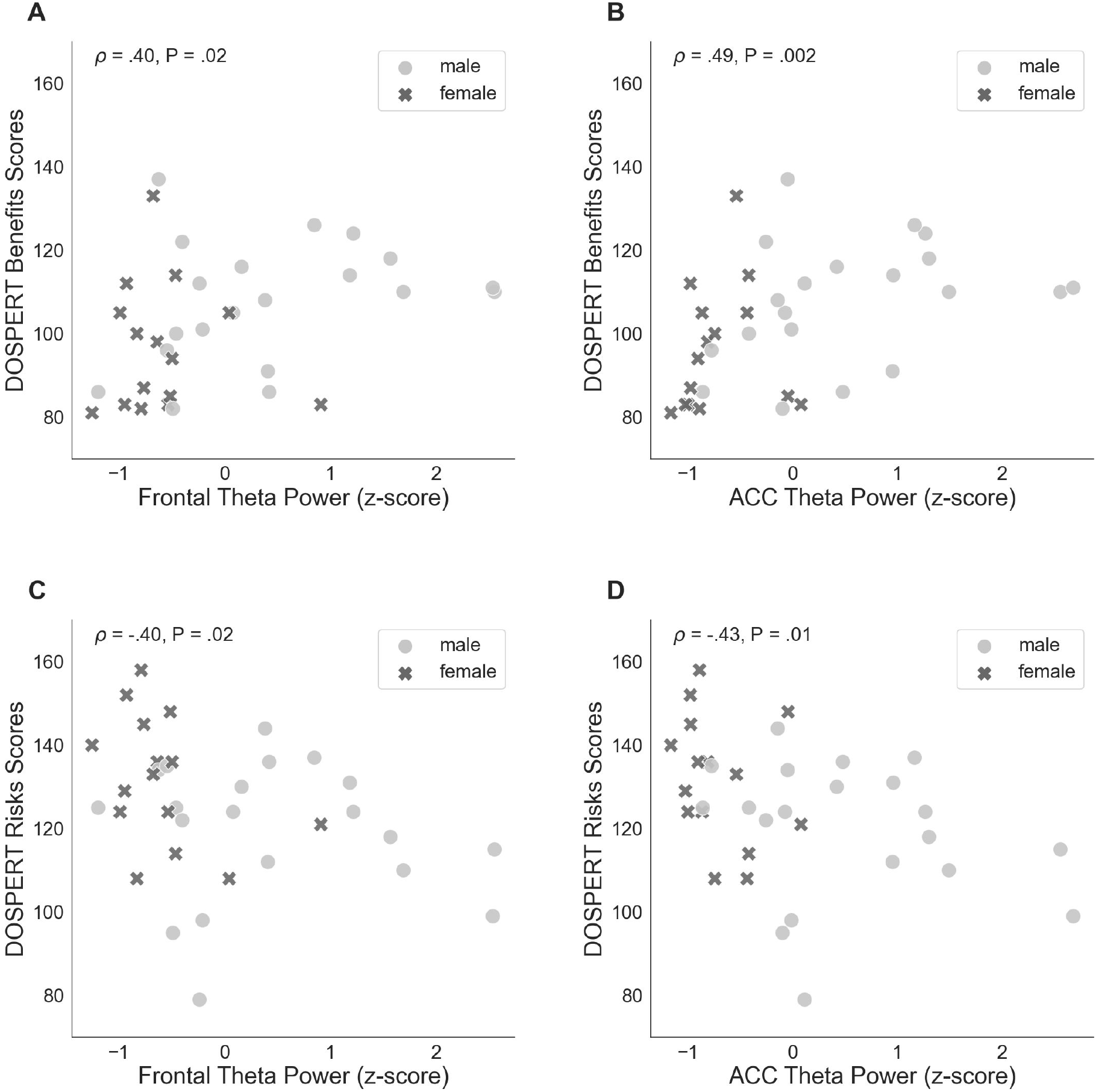
Spearman’s correlations of self-assessed measures of reward (DOSPERT benefits) and punishment (DOSPERT risks) sensitivity with frontal theta power and ACC theta power, standardized. DOSPERT - Domain-Specific Risk-Taking Scale. ACC - anterior cingulate cortex. P - unadjusted p-value that was significant after controlling for the false discovery rate at level q = 0.05.

### 3.4 ACC Theta Power

ACC theta power strongly correlated with rsFT in sensor space, as expected (Spearman’s ρ = 0.9, P = 1.34e-13). Next, we observed a significant non-parametric association between theta power in the ACC and average risk taking (Spearman’s ρ = 0.49, P = 0.003, FDR-controlled), as well as first-trial risk taking (Spearman’s ρ = 0.51, P = 0.002, FDR-controlled) (Figure 6). Moreover, ACC theta power significantly correlated with DOSPERT benefits (Spearman’s ρ = 0.49, P = 0.002, FDR-controlled) and DOSPERT risks (Spearman’s ρ = −0.43, P = 0.01, FDR-controlled) subscales (Figure 5). ACC theta power did not correlate with DOSPERT likelihood scores. Finally, males had higher ACC theta power than females (Mann-Whitney U-test P = 0.000006). In sum, the results obtained for the ACC theta power were similar to the results obtained for the rsFT - yet the former were more pronounced as reflected in higher Spearman’s ρ values, which was also replicated based on the resting-state recording after the game.

**Figure 6.**
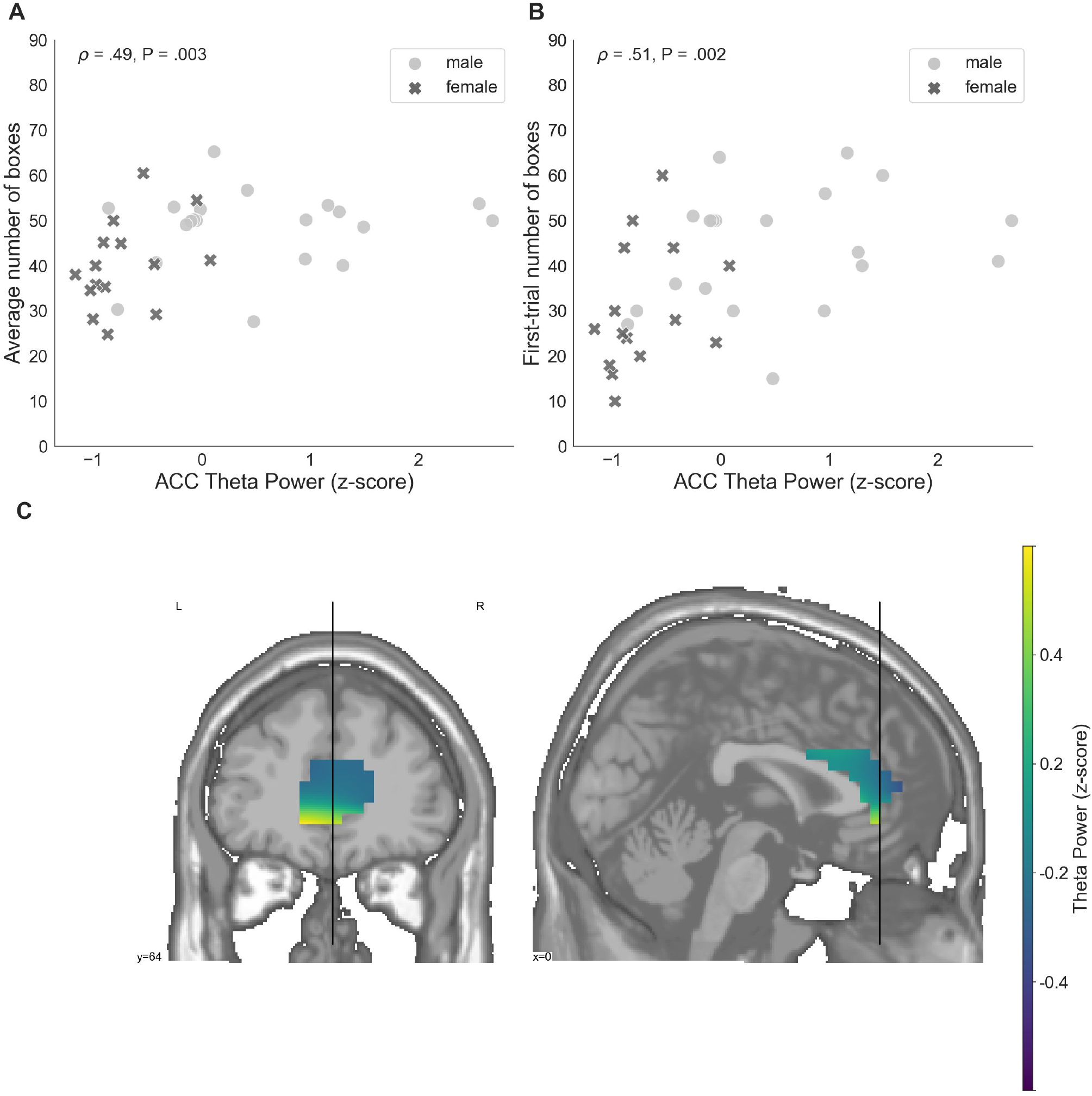
(**A**) A relation between the average number of selected boxes during the game and average theta power in voxels located in the ACC, standardized. (**B**) The number of boxes selected in the first trial of the game and average theta power in voxels localized to the ACC, standardized. (**C**) Theta power in voxels localized to the ACC, standardized. ACC - anterior cingulate cortex. P - unadjusted p-value that was significant after controlling the false discovery rate at level q = 0.05.

### 3.5 Mediation

Post-hoc mediation analysis revealed that the effect of the ACC theta power on average risk taking in the game was fully mediated by the sensitivity to rewards - DOSPERT benefits scores (Figure 7). The indirect effect (ACME) was statistically significant (P = 0.03): β = 1.5, CI = [0.11 - 3.34]. At the same time, average direct effect (ADE) was insignificant, indicating the complete mediation. Further mediation analysis demonstrated that the effect of the ACC theta power on first-trial risk taking was partially mediated by the DOSPERT benefits subscale. The indirect effect (ACME) was statistically significant (P = 0.002): β = 2.9, CI = [0.9 - 5.68]. However, average direct effect (ADE) was also significant (P = 0.02), indicating the incomplete mediation. Moreover, being motivated by the idea of the present study, we further suggested that sex differences in risk taking may be mediated by reward sensitivity relating to resting-state theta oscillations in the ACC. To formally test this suggestion, we extended the mediation model by allowing sequential meditation. The results partially confirmed this hypothesis: structural equation modeling (SEM) revealed that the indirect pathway of the effect of sex on first-trial risk taking via the ACC theta power and DOSPERT benefits scores was significant (P = 0.03): β = 3.7, CI = [0.83 - 7.3]. Moreover, it fully accounted for the overall impact of sex on first-trial risk taking with the direct effect being insignificant. However, we found no significant sequential mediation with respect to the average risk taking. Details of SEM models are presented in Supplementary Figures 7 and 8.

**Figure 7.**
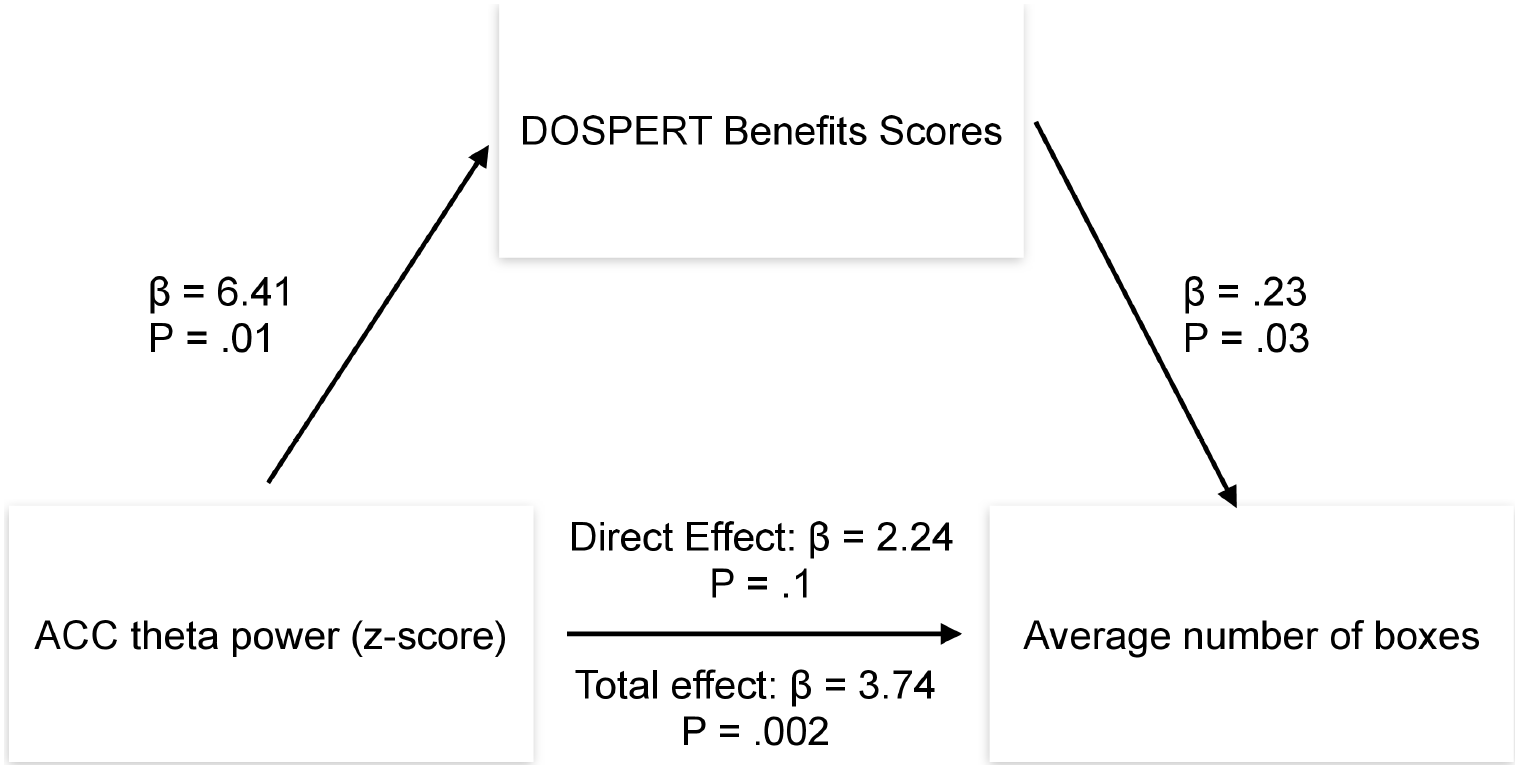
Results of the mediation analysis. β - regression coefficient; P - p-value of a regression coefficient. DOSPERT - Domain-Specific Risk-Taking Scale. ACC - anterior cingulate cortex.

### 3.6 Regression Analysis

Regression analysis demonstrated that the interaction term of sex with rsFTA in relation to risk-taking measures was not significant (as measured by ANOVA; p-values of the F-test for comparing models with and without the interaction term were 0.12 and 0.33 respectively for average and first-trial risk taking) i.e., sex did not modulate linear relationship of rsFTA and risk-taking. However, an analogous regression analysis based on resting-state recordings after the game revealed that sex significantly interacted with rsFTA after the game in relation to average risk-taking in the game (the p-value of the F-test for comparing models with and without the interaction term was 0.04). Importantly, inclusion of sex as a control variable in linear models where rsFTA was the independent variable significantly improved performance of models (F-test p-values were 0.03 and 0.01 respectively for average and first-trial risk taking). Furthermore, it marginally improved performance of a model for effect of rsFT on average (F-test P = 0.05), but not first-trial risk taking (F-test P = 0.09). At the same time, inclusion of sex as a control variable in linear models where ACC theta power was the independent variable did not significantly improve performance of models (F-test p-values were 0.16 and 0.20 respectively for average and first-trial risk taking). Finally, regression of average risk taking on ACC theta power before the game was significantly improved by controlling for rsFTA (F-test P = 0.03), while it was not improved based on the resting-state activity after the game (F-test P = 0.15). Regression results are provided in Supplementary Tables 2 and 3.

## 4 Discussion

Using resting-state MEG recordings and three distinct measures of risk attitudes, we show that sex differences in risk taking are associated with reward sensitivity, which, in turn, are linked to resting-state ACC theta oscillations (Figure 8). On the behavioral level, males were more sensitive to rewards than females. Game-based measures of risk taking showed significant sex differences and also correlated with self-reported expected benefits of risky actions (DOSPERT benefits scores). On the neural level, rsFTA explained average risk taking during the repeated game exclusively in the female subsample. By contrast, in the whole sample, rsFT correlated with first-trial risk taking and also with DOSPERT benefits and risks scores, indicating an association with reward and punishment sensitivity. Finally, due to a refined spatial specificity, theta power localized to the ACC correlated with outcome sensitivities and game-based measures of risk taking even more strongly than the rsFT did. The findings suggest that resting-state ACC activity is a possible source of sex differences in reward sensitivity, and, consequently, in risk taking.

**Figure 8.**
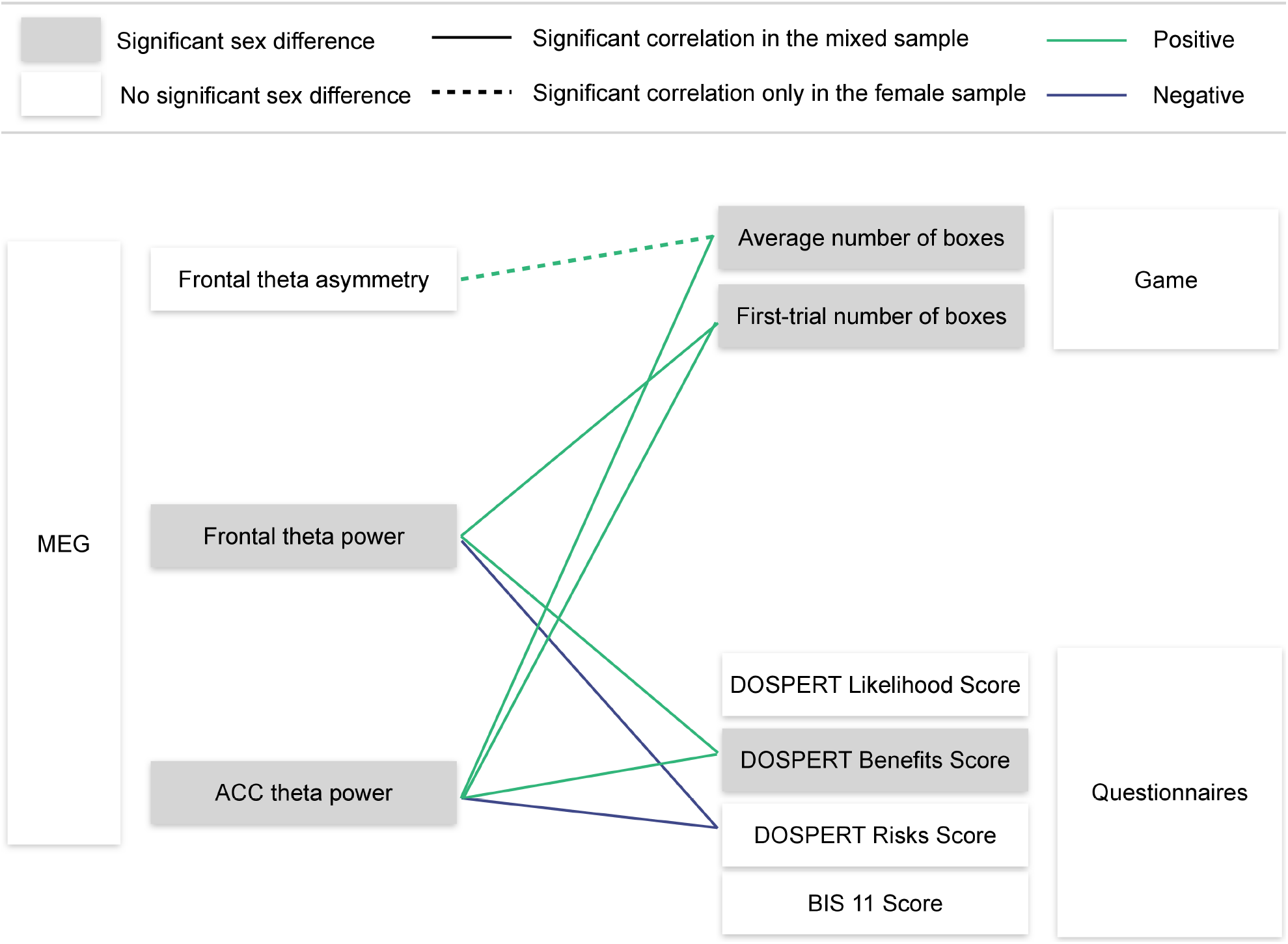
Visualization of the main findings. MEG - magnetoencephalography. ACC - anterior cingulate cortex. DOSPERT - Domain-Specific Risk-Taking Scale. BIS11 - Barratt Impulsiveness Scale. Significant correlations are reported after FDR adjustment.

### 4.1 Behavioral Measures

DOSPERT benefits subscale significantly correlated with both average and first-trial risk taking, converging with previous studies (Weber, Blais, & Betz, 2002; Hanoch, Johnson, & Wilke, 2006; Fukunaga et al., 2018). Further, self-assessed reward sensitivity demonstrated a greater correlation with first-trial than with average risk taking, indicating that sensitivity to outcomes could affect the former more (Erev, Ert, & Yechiam, 2008; Lejarranga & Gonzalez, 2011). Absence of correlations between BIS11 scores and performance on decision-making tasks is in line with the literature (Lauriola, Panno, Levin, & Lejuez, 2014; Reddy et al., 2014; Hüpen, Habel, Schneider, Kable, & Wagels, 2019; Gomide Vasconcelos, Sergeant, Corrêa, Mattos, & Malloy-Diniz, 2014).

We observed significant sex differences in first-trial and average risk taking during the game, which was expected based on the extensive literature on sex differences in decision-making under uncertainty (e.g., Jianakoplos & Bernasek, 1998; Weber et al., 2002; Charness & Gneezy, 2012; Zhou et al., 2014). Notably, however, Crosetto & Filippin (2013) did not report significant sex differences in their versions of BRET. This discrepancy could be explained by the use of repeated trials or more salient financial incentives in our task. Furthermore, it has been observed earlier that some measures reveal that females are more risk-averse than males, while others do not (e.g., Charness, Gneezy, & Imas, 2013; Filippin & Crosetto, 2016). Our finding that there was no significant sex difference in DOSPERT likelihood scores supports this observation.

As for the reward and punishment sensitivities, we observed that on average males scored higher than females on DOSPERT benefits subscale. Previous studies also reported sex differences in outcome sensitivities based on DOSPERT (Weber et al., 2002; Hanoch et al., 2006; Lee & Jeong, 2013) and other measures (Li, Huang, Lin, & Sun, 2007; Cross, Copping, & Campbell, 2011). In line with previous studies, we found no significant sex differences in impulsivity (Kamarajan et al., 2008; Lee & Jeong, 2013; Liu, Zubieta, & Heitzeg, 2012). Summarizing the facts presented above, behavioral evidence from the current work suggests that sensitivity to outcomes, rather than impulsivity, is a candidate trait that could explain sex differences in risk attitudes.

### 4.2 Frontal Theta Asymmetry (rsFTA)

Analysis of the MEG oscillatory activity showed no significant sex differences in rsFTA, converging with previous EEG work (Ocklenburg et al., 2019). Therefore, as predicted, we simultaneously observed, (1) sex differences in risk taking based on game performance, and (2) no sex differences in the neural trait previously associated with this decision-making characteristic.

We found a significant positive correlation of rsFTA with average risk taking exclusively in the female subsample, confirming earlier findings in female populations by Gianotti et al. (2009). Average and first-trial risk taking in the game did not correlate with rsFTA in the joint sample, which is in contrast with the result of Studer et al. (2013). However, Studer et al. (2013) did not report sex-specific results, and this study contained 70% of females, which, according to our findings, could bias the result obtained for the joint sample. Because regression analyses demonstrated significant effects of rsFTA on average risk taking in the game and because we observed correlation coefficients of rsFTA with measures of risk taking (although insignificant) comparable in magnitude to previous work with larger samples (Studer et al., 2013), a possible interpretation is that we were not able to reliably detect significant non-parametric associations between rsFTA and risk taking for the whole sample due to the limited sample size.

Higher rsFTA may be associated with the lower relative right frontal activity, and thus the prevalence of left frontal activity (Gianotti et al., 2009, Studer et al., 2013), which is partially supported by previously observed negative associations between theta power and cortical activity (Oakes et al., 2004; Scheeringa et al., 2008). Additional evidence that frontal lateralization is related to risk taking comes from stimulation studies, focused on the role of right DLPFC in decision-making (Knoch et al., 2006; Fecteau et al., 2007a; Fecteau et al., 2007b; Cho et al., 2010; Sela, Kilim, & Lavidor, 2012). Furthermore, several studies reported sex differences in the involvement of the right and left frontal cortices in decision-making (Bolla, Eldreth, Matochik, & Cadet, 2004; Tranel, Damasio, Denburg, & Bechara, 2005; Neo & McNaughton, 2011). Our findings contribute to the evidence that sex may interact with frontal asymmetry in relation to risk taking, but this requires further testing.

### 4.3 Theta Power

We found a strong association between rsFT and theta power in the ACC. This outcome is consistent with previous dipole-fitting studies that revealed possible sources of rsFT in the ACC (Asada, Fukuda, Tsunoda, Yamaguchi, & Tonoike, 1999; Scheeringa et al., 2008; Clemens et al., 2010).

#### 4.3.1 Risk Taking

We report on the existence of the significant positive correlation between rsFT and first-trial risk taking. Two previous studies did not find correlations of rsFT with risk taking (Massar et al., 2012; Studer et al., 2013). There are three notable similarities between their experimental designs that differentiate them from our paradigm. First, both studies introduced losses in the task either explicitly or via a safe gamble (Ert & Erev, 2013). Second, they forced participants to choose between two gambles with the same expected value (Massar et al., 2012) or with very similar expected values (Studer et al., 2013). Third, the computation of expected values of gambles in tasks used by Massar et al. (2012) and Studer et al. (2013) was straightforward. Therefore, differences in experimental designs associated with values and presentation of options might have affected the observed correlations between rsFT and risk taking.

All observed correlations for rsFT were even stronger for the ACC theta power, and it also significantly correlated with average risk taking in the game. It is an expected result. If rsFT originates at the level of the ACC (Asada et al., 1999; Scheeringa et al., 2008; Clemens et al., 2010), then the results would be more pronounced at the source level compared to the sensor level due to the contamination of sensor-level activity from other less relevant sources. Thus, our findings are aligned with the extensive neuroimaging research demonstrating the involvement of the ACC in decisions under risk (Paulus & Frank, 2006; Christopoulos et al., 2009; Hewig et al., 2009; Mohr, Biele, & Heekeren, 2010; Schonberg et al., 2012; Fukunaga et al., 2018).

#### 4.3.2 Reward and Punishment Sensitivity

Additionally, we found strong correlations of self-assessed punishment (DOSPERT risks) and reward (DOSPERT benefits) sensitivities with rsFT and the ACC theta power. Few previous studies examined associations between rsFT or ACC activity at rest and outcome sensitivity. Our findings contribute to the evidence that rsFT is related to outcome sensitivity (Massar et al., 2012; Massar et al., 2014). Regarding theta oscillations and ACC activity during tasks, both measures have previously been associated with reactions to rewards and punishments (Debener et al., 2005; Cohen et al., 2007; Kamarajan et al., 2008; Santesso et al., 2011; Crowley et al., 2014; Van Duijvenvoorde et al., 2014). Research in humans (Gehring & Willoughby, 2002; Wang, Ulbert, Schomer, Marinkovic, & Halgren, 2005; Iannaccone et al., 2015) and primates (Tsujimoto, Shimazu, Isomura, & Sasaki, 2010; Womelsdorf, Johnston, Vinck, & Everling, 2010; Babapoor-Farrokhran, Vinck, Womelsdorf, & Everling, 2017; Taub, Perets, Kahana, & Paz, 2018) singled out theta ACC activity as a source of signals associated with feedback and behavioral adjustment. Our results further extend this literature.

#### 4.3.3 Sex Differences

We found significant sex differences in rsFT. However, evidence from previous studies is mixed. Zappasodi et al. (2006) also used MEG and reported the presence of sex differences in resting-state theta power. Other studies used EEG and reported no significant sex differences (Jausovec & Jausovec, 2010; Gmehlin et al., 2011; Kober & Neuper, 2011; Banis, Geerligs, & Lorist, 2014), or higher theta power in females compared to males (Clarke, Barry, McCarthy, & Selikowitz, 2001; Kamarajan et al., 2008; Osinsky et al., 2017). We examined the demographic characteristics of participants in these studies and did not find a pattern that could account for such inconsistent results. One possibility is that sex difference in skull conductivities affects EEG recordings but not MEG (Huttunen, Wikström, Salonen, & Ilmoniemi, 1999).

Sex differences in the resting-state ACC theta power were even more pronounced. It is in line with the diverse evidence from previous studies that demonstrated sex differences associated with this region (Goldstein et al., 2001; Markham & Juraska, 2002; Zhou et al., 2014). These sex differences may be linked to levels of testosterone and its effects on midbrain dopaminergic pathways (Johnson et al., 2010). On the one hand, activity of the ACC is associated with dopaminergic projections from the midbrain (Holroyd & Coles, 2002), and dopaminergic genetic polymorphisms correlate with risk taking and also with amplitudes of FRN (Heitland et al., 2012). On the other hand, higher levels of testosterone are associated with risk taking (Apicella et al., 2008; Stanton, Liening, & Schultheiss, 2011) and also with outcome sensitivity (Van Honk et al., 2004). Therefore, baseline ACC activity may be linked to sex differences in outcome sensitivity and, consequently, risk taking. It should be noted, however, that the previous literature on sex differences in either rsFT or resting ACC theta activity is rather scarce. Accordingly, validation of the current results in future MEG and combined MRI-EEG studies will be necessary.

Noticeably, a mediation analysis allowed us to formally test the hypothesis that sex differences in risk taking may be mediated by reward sensitivity via resting-state theta oscillations in the ACC. The results revealed that reward sensitivity assessed via DOSPERT benefits scores mediated the effects of resting-state ACC theta oscillations on average and first-trial risk taking in the game. Furthermore, structural equation modeling demonstrated that the indirect pathway of the effect of sex on first-trial risk taking via the ACC theta power and DOSPERT benefits scores was significant; it fully accounted for the overall impact of sex on first-trial risk taking. Therefore, even though there may be other confounding variables, reward sensitivity is a candidate trait for explaining sex differences in risk taking where resting-state ACC activity is a potential contributing mechanism.

Finally, regression analysis demonstrated that rsFTA and sex captured significantly different portions of variance in task performance, while ACC theta power explained variance due to sex. Therefore, if we only had information about rsFTA we would not be able to explain variability in risk taking of participants associated with their sex, while this can be done based on resting ACC theta power. Interestingly, Weis et al. (2020) has recently shown that sex classification based on resting-state connectivity of ACC can be done with 74.4% accuracy. In addition, results of regression analysis showed that average performance in the game was explained best when including both rsFTA and ACC theta power before the game (as opposed to including only one of these characteristics) which further highlights a possibility for functionally distinct involvement of these neural traits in risk-taking. Future research is required to clarify this question.

### 4.4 Limitations

This study has several limitations. An important drawback of our experiment is not controlling for the menstrual cycle phase of female participants. This could have confounded our results because cortical activity is affected by menstrual cycle phase and blood estrogen level (Dietrich et al., 2001; Hausmann, 2005). Furthermore, we had a relatively small sample size (35), a higher number of males compared to females (20/15) and a rather young group of participants. Due to this limitation, we could not reliably detect correlations of risk-taking measures with neural traits in male and female subsamples separately (although technically it is possible). Thus, we were mostly interpreting results relating to the joint sample as a whole. Our findings necessitate further research with larger samples, separately for males and females. Nevertheless, power analysis suggests that our joint sample was sufficient to detect correlation coefficients of 0.45 or higher. Reliability of our results was further confirmed by replicating significant results based on resting-state recordings after the game. Another limitation is that we did not have individual MRI of participants which could have improved our source-modeling results even further. Finally, it must be noted that the study was correlational, and thus we could not establish direct causal links - this critique, however, applies to almost all EEG/MEG studies.

### 4.5 Conclusions

Our study provides novel evidence for the understanding of sex-related differences in risk taking by suggesting that these differences arise due to lower reward sensitivity in females as compared to males. Further, these differences are associated with resting-state theta band activity in the ACC. In addition, we find evidence that sex interacts with neural traits in relation to risk taking. Thus, our results stress the necessity to control for sex in decision neuroscience studies, as also suggested earlier (Cahill, 2006). Overall, we provide evidence that different measures of risk taking are differentially associated with distinct neural traits. This in turn suggests that various risk-preference elicitation methods may involve several ‘cognitive strategies’ (Pedroni et al., 2017). This could be the reason why some measures of risk taking demonstrate sex differences while others do not. Our findings indicate that when sex differences according to a specific risk-taking measure are pronounced, the ACC theta power significantly correlates with risk taking in the sample containing both males and females. Finally, our results contribute to a broader topic of sex differences in decision-making and its dysfunction. In fact, differences in reward processing may be involved in more prevalent rates of obesity, anxiety and depression among females (Loxton & Dawe, 2007; Cavanagh et al., 2019).

## 6 Author Contributions

Maria Azanova: Conceptualization, Methodology, Software, Investigation, Data Curation, Formal analysis, Writing - Original Draft, Visualization. Maria Herrojo Ruiz: Formal analysis, Writing - Review & Editing, Visualization. Alexis V. Belianin: Methodology, Writing - Review & Editing. Vasily Klucharev: Writing - Review & Editing, Funding acquisition. Vadim V. Nikulin: Methodology, Writing - Review & Editing, Supervision.

## 8 Abbreviations

rsFTA: resting-state frontal theta asymmetry (right-left)
rsFT: resting-state frontal theta power
DOSPERT: Domain-Specific Risk-Taking scale
MEG: magnetoencephalography
EEG: electroencephalography
fMRI: functional magnetic resonance imaging
ACC: anterior cingulate cortex
BRET: Bomb Risk Elicitation Task
BIS11: Barratt Impulsiveness Scale

## 9 Acknowledgments

Supported by the International Laboratory of Social Neurobiology ICN HSE RF Government grant ag. No. 075-15-2019-1930. MA is supported by the German Federal Ministry of Education and Research (BMBF) and the Max Planck Society.

## 10 Data Availability Statement

The datasets are available from corresponding authors upon a reasonable request.

