## Supplementary Material for "Resting-state theta oscillations and reward sensitivity in risk taking"

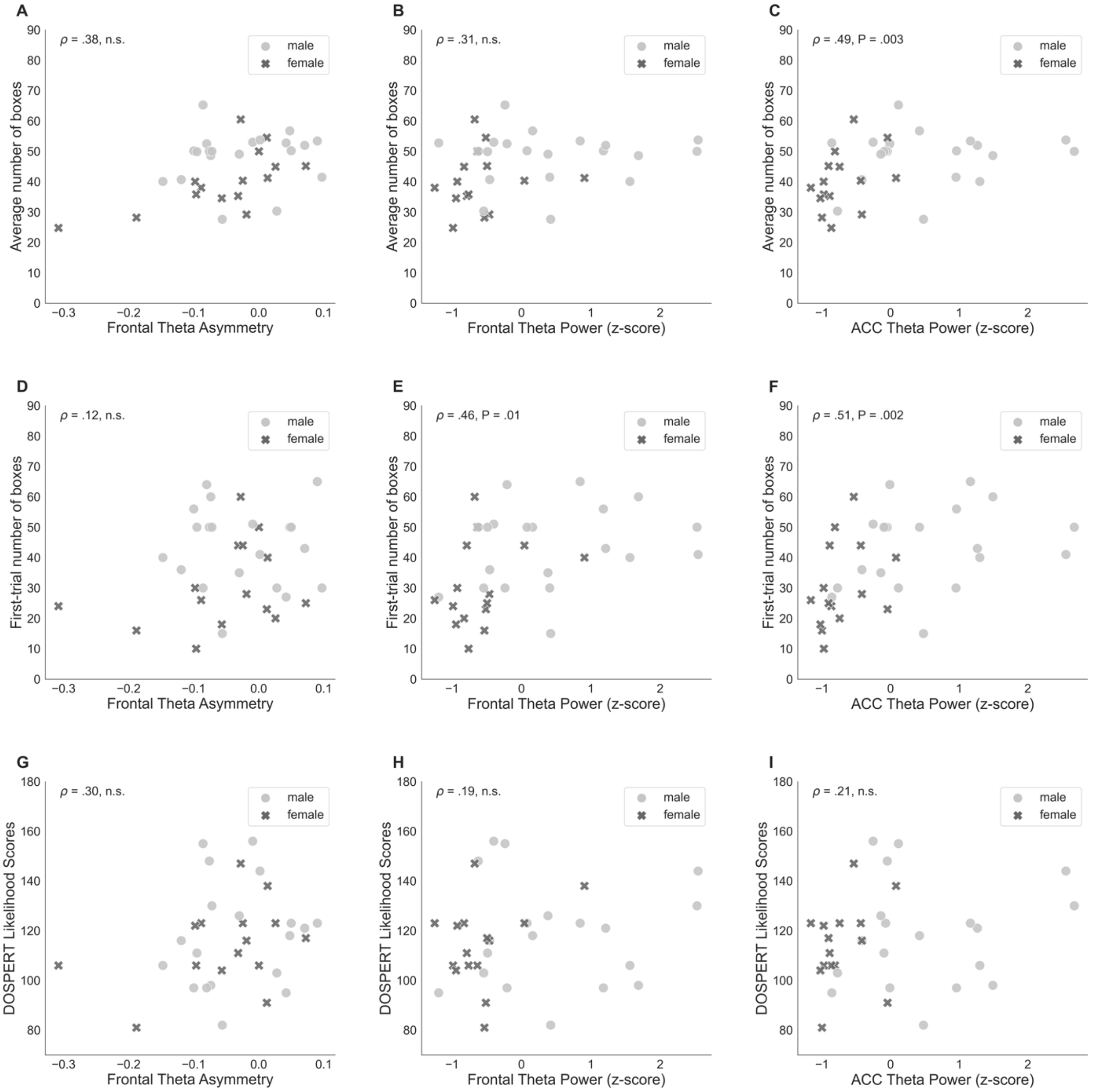


**Supplementary Figure 1.** Spearman's correlations of risk-taking measures and neurobiological measures based on resting-state recordings before the game. P-values are reported only for correlations significant after FDR-control. DOSPERT - Domain-Specific Risk-Taking Scale. ACC - anterior cingulate cortex.

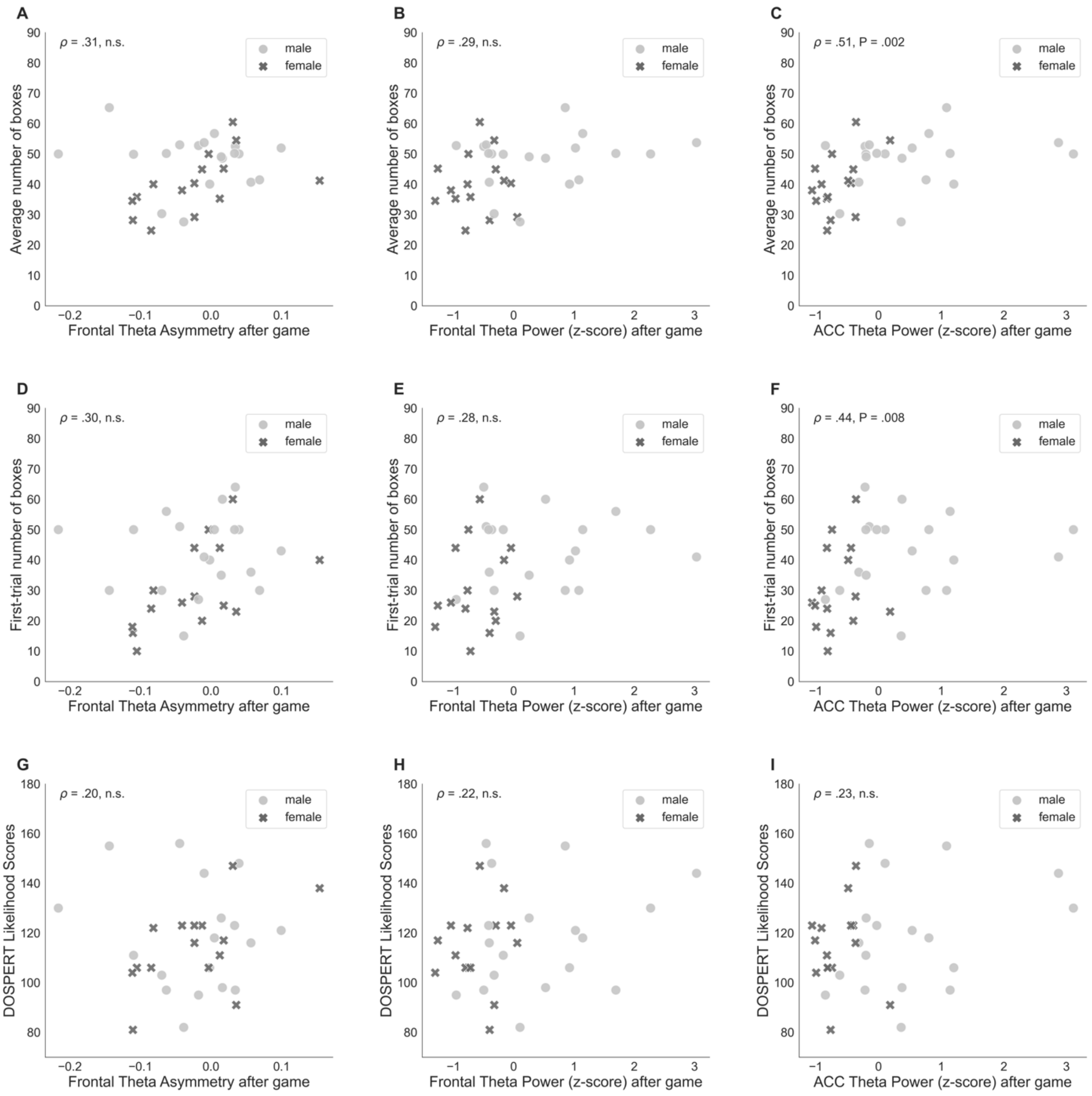


**Supplementary Figure 2.** Spearman's correlations of risk-taking measures and neurobiological measures based on resting-state recordings after the game. P-values are reported only for correlations significant after FDR-control. DOSPERT - Domain-Specific Risk-Taking Scale. ACC - anterior cingulate cortex. Females and males did not differ significantly in rsFTA after the game (Mann-Whitney U-test P = 0.47). Males had significantly higher rsFT (Mann-Whitney U-test P = 0.0006) and ACC theta power (Mann-Whitney U-test P = 7.68e-06) after the game than females. rsFTA before and after the game showed a significant, but moderate non-parametric association (Spearman's $\rho = 0.47$, P = 0.004, FDR-controlled), while rsFT before and after the game correlated more strongly (Spearman's $\rho = 0.85$, P = 2.73e-10, FDR-controlled). Finally, ACC theta power before and after the game highly and significantly correlated as well (Spearman's $\rho = 0.94$, P = 1.19e-16, FDR-controlled).

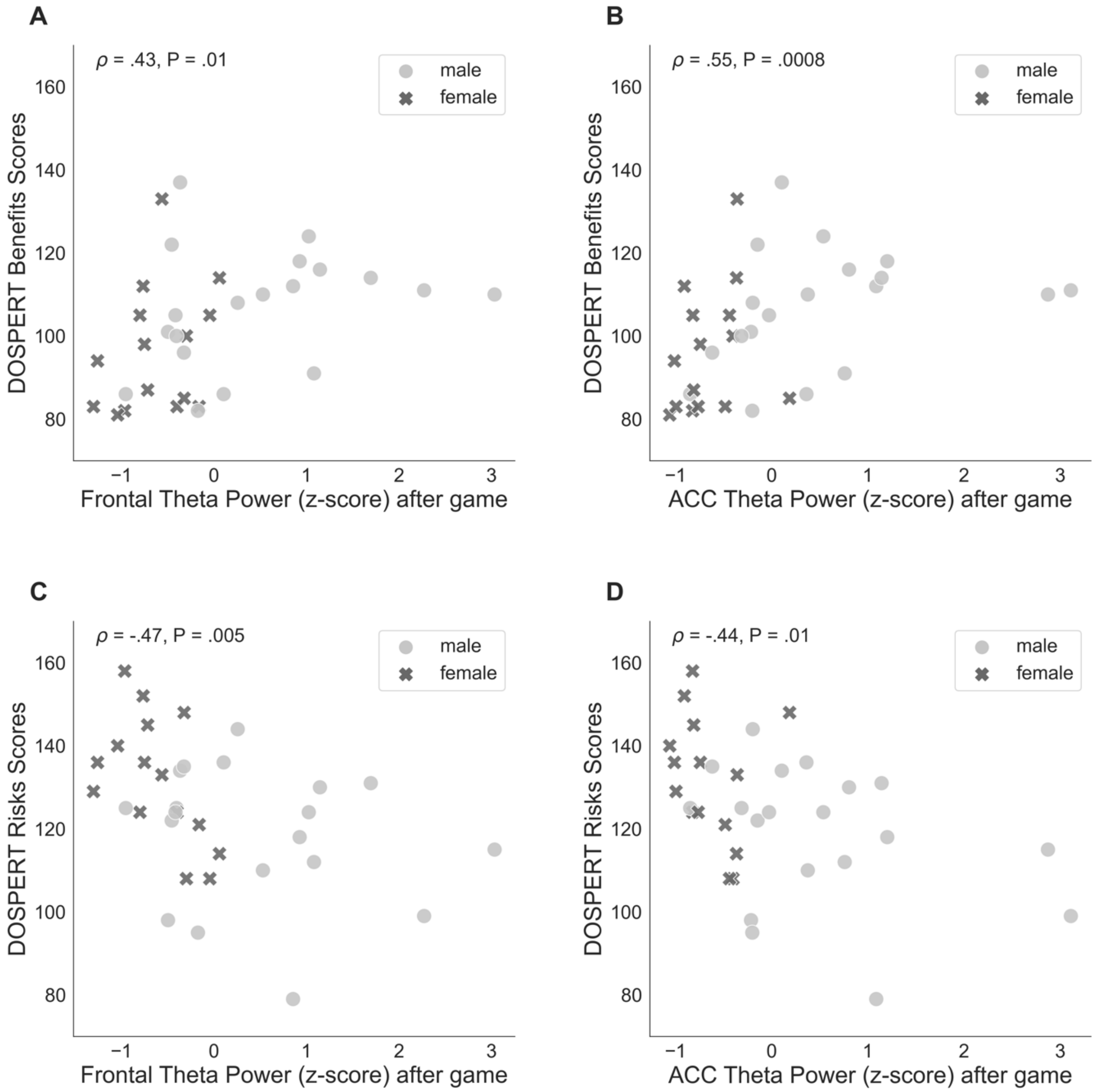


**Supplementary Figure 3.** Spearman's correlations of self-assessed measures of reward (DOSPERT benefits) and punishment (DOSPERT risks) sensitivities with frontal theta power and ACC theta power, standardized. P-values are reported only for correlations significant after FDR-control. DOSPERT - Domain-Specific Risk-Taking Scale. ACC - anterior cingulate cortex.

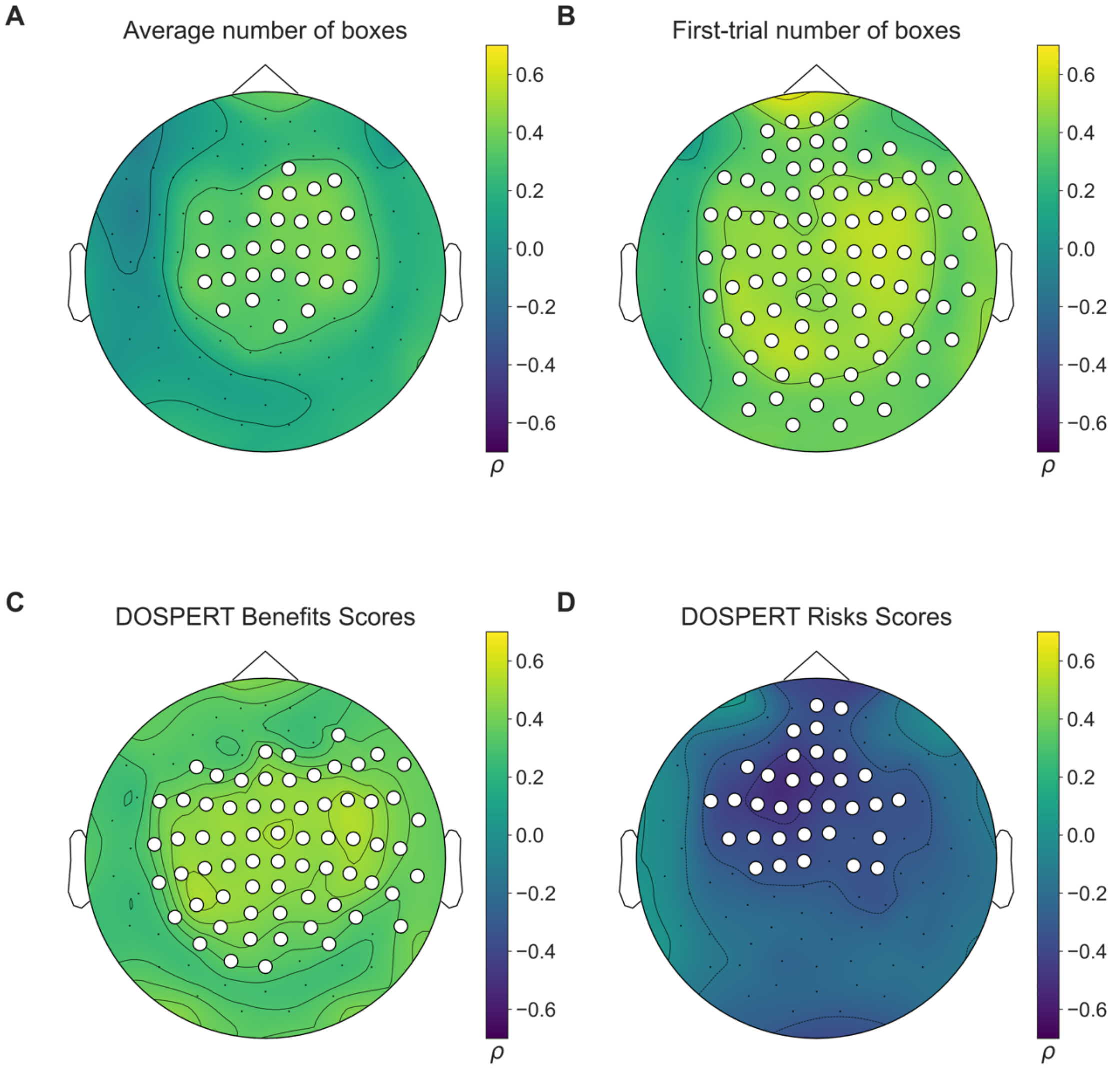


**Supplementary Figure 4.** Results of non-parametric statistical clustering in sensor space (1024 permutations) based on resting-state recordings before the game. Color denotes Spearman's correlation coefficients of theta power at sensor locations (102 magnetometers) with (A) average number of boxes chosen in the game; (B) number of boxes chosen in the first trial; (C) DOSPERT expected benefits subscale scores; and (D) DOSPERT perceived risks subscale scores. White dot markers denote magnetometers included in a statistically significant cluster (alpha level of 0.05). DOSPERT - Domain-Specific Risk-Taking Scale.

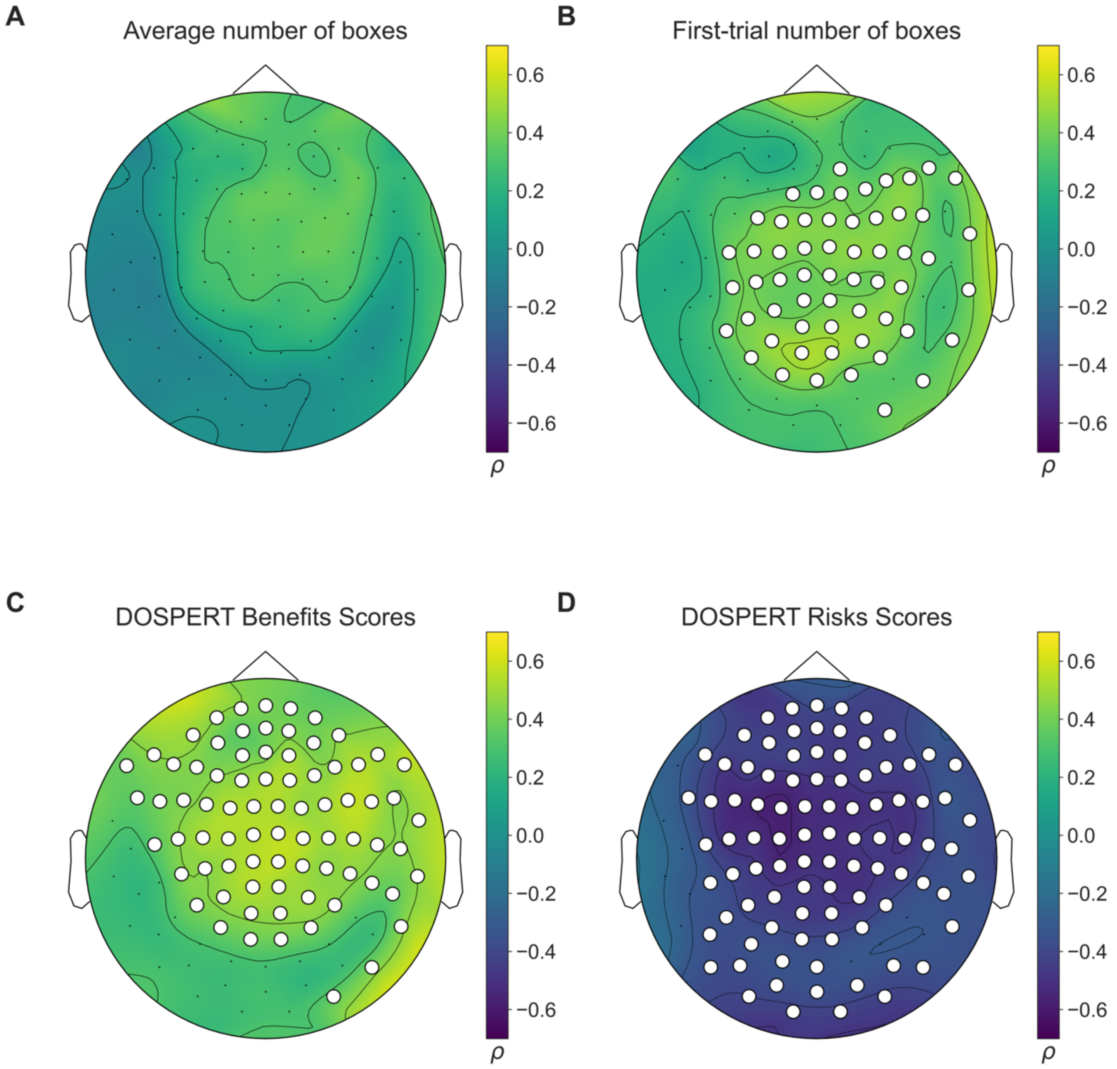


**Supplementary Figure 5.** Results of non-parametric statistical clustering in sensor space (1024 permutations) based on resting-state recordings after the game. Color denotes Spearman's correlation coefficients of theta power at sensor locations (102 magnetometers) with (A) average number of boxes chosen in the game; (B) number of boxes chosen in the first trial; (C) DOSPERT expected benefits subscale scores; and (D) DOSPERT perceived risks subscale scores. White dot markers denote magnetometers included in a statistically significant cluster (alpha level of 0.05). DOSPERT - Domain-Specific Risk-Taking Scale.

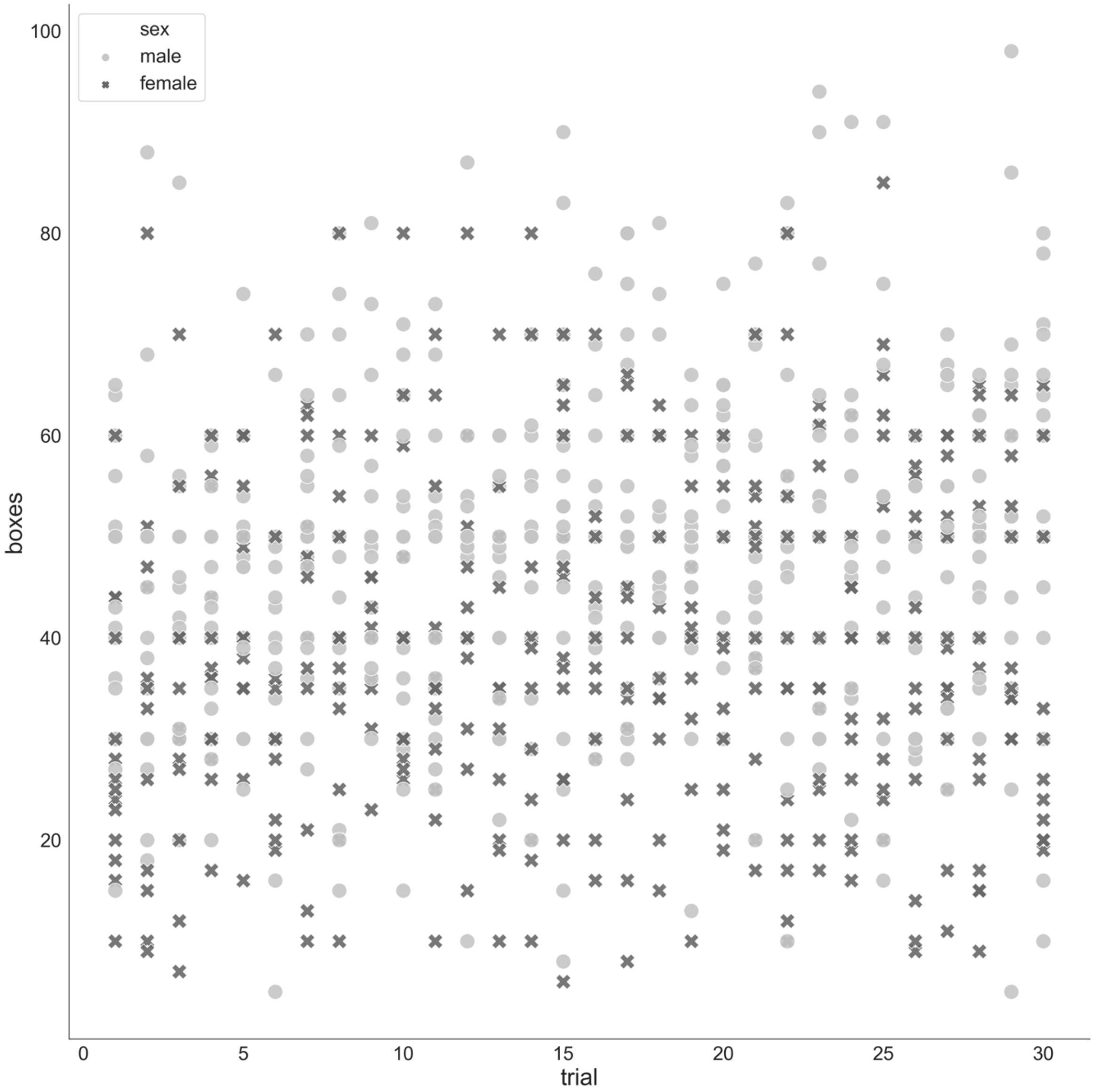


**Supplementary Figure 6.** Numbers of boxes chosen by each participant in each trial (n = 1041, trials with 0, 1 and 2 chosen boxes were excluded).

| | rsFT | ACC theta | rsFT after game | ACC theta after game |
|---|---|---|---|---|
| % change after loss | 0.33 (n.s.) | 0.41 (0.01) | 0.41 (0.02) | 0.49 (0.003) |
| % change after win | -0.33 (n.s.) | -0.43 (0.01) | -0.29 (n.s.) | -0.47 (0.005) |
| Difference % change after win and loss | -0.37 (0.03) | -0.49 (0.002) | -0.36 (0.03) | -0.54 (0.001) |

**Supplementary Table 1.** Spearman's correlation coefficients of rsFT/ACC theta power and average percent changes in the number of boxes after losing or winning. P-values for correlations significant after FDR-control are reported in parentheses.

Both average percent change after win (mean = 0.26; SD = 0.14) and percent change after loss (mean = -0.08; SD = 0.14) measures were statistically different from 0 (t-test p-values were 9.42e-07 and 0.001 respectively). We observed significant sex differences in percent changes in the numbers of boxes chosen after losing (Mann-Whitney U-test P = 0.007; mean males = -0.03; mean females = -0.15) and winning (Mann-Whitney U-test P = 0.01; mean males = 0.18; mean females = 0.36), and in the difference of these reactions (Mann-Whitney U-test P = 0.003; mean males = 0.21; mean females = 0.51).

| | *Dependent variable:* | | | | | | | |
|---|---|---|---|---|---|---|---|---|
| | Average number of boxes | | | | First-trial number of boxes | | | |
| | (1) | (2) | (3) | (4) | (5) | (6) | (7) | (8) |
| rsFTA before | 41.19** | 67.23*** | | | 20.01 | 45.84 | | |
| | (17.94) | (23.79) | | | (28.56) | (38.83) | | |
| rsFTA after | | | 20.43 | 74.77** | | | 48.05 | 111.73** |
| | | | (21.91) | (33.02) | | | (30.66) | (47.22) |
| male | 7.03** | 4.82 | 7.84** | 5.93* | 12.55** | 10.35* | 11.76** | 9.52** |
| | (2.99) | (3.22) | (3.20) | (3.17) | (4.76) | (5.26) | (4.48) | (4.53) |
| rsFTA before:male | | −56.81 | | | | −56.35 | | |
| | | (35.13) | | | | (57.35) | | |
| rsFTA after:male | | | | −89.95** | | | | −105.43* |
| | | | | (42.48) | | | | (60.76) |
| Constant | 42.42*** | 43.83*** | 40.65*** | 41.91*** | 31.62*** | 33.02*** | 31.64*** | 33.12*** |
| | (2.43) | (2.53) | (2.45) | (2.40) | (3.87) | (4.12) | (3.43) | (3.43) |
| Observations | 35 | 35 | 34 | 34 | 35 | 35 | 34 | 34 |
| $R^2$ | 0.29 | 0.34 | 0.18 | 0.29 | 0.21 | 0.23 | 0.24 | 0.31 |
| Adjusted $R^2$ | 0.24 | 0.28 | 0.13 | 0.22 | 0.16 | 0.16 | 0.19 | 0.24 |
| *Note:* | | | | | | *p<0.1; **p<0.05; ***p<0.01 | | |

**Supplementary Table 2.** Regression results used to assess interactions of rsFTA before and after the game and sex in relation to game-based risk-taking measures.

| | *Dependent variable:* | | | | | | | |
|---|---|---|---|---|---|---|---|---|
| | Average number of boxes | | | | First-trial number of boxes | | | |
| | (1) | (2) | (3) | (4) | (5) | (6) | (7) | (8) |
| ACC theta before | 3.75** | 3.16** | | | 6.79*** | 6.51*** | | |
| | (1.60) | (1.52) | | | (2.32) | (2.37) | | |
| rsFTA before | | 42.07** | | | | 19.83 | | |
| | | (18.24) | | | | (28.33) | | |
| ACC theta after | | | 4.15** | 4.52*** | | | 4.84* | 5.56** |
| | | | (1.60) | (1.59) | | | (2.40) | (2.32) |
| rsFTA after | | | | 31.81 | | | | 62.54* |
| | | | | (21.57) | | | | (31.51) |
| Constant | 44.86*** | 46.47*** | 44.61*** | 45.26*** | 38.03*** | 38.78*** | 37.24*** | 38.51*** |
| | (1.58) | (1.64) | (1.58) | (1.61) | (2.28) | (2.54) | (2.36) | (2.35) |
| Observations | 35 | 35 | 34 | 34 | 35 | 35 | 34 | 34 |
| $R^2$ | 0.14 | 0.27 | 0.17 | 0.23 | 0.21 | 0.22 | 0.11 | 0.21 |
| Adjusted $R^2$ | 0.12 | 0.22 | 0.15 | 0.18 | 0.18 | 0.17 | 0.09 | 0.16 |
| *Note:* | | | | | | | | $^{*}$p<0.1; $^{**}$p<0.05; $^{***}$p<0.01 |

**Supplementary Table 3.** Regression results for linear models of risk taking on the two measures of neural characteristics.

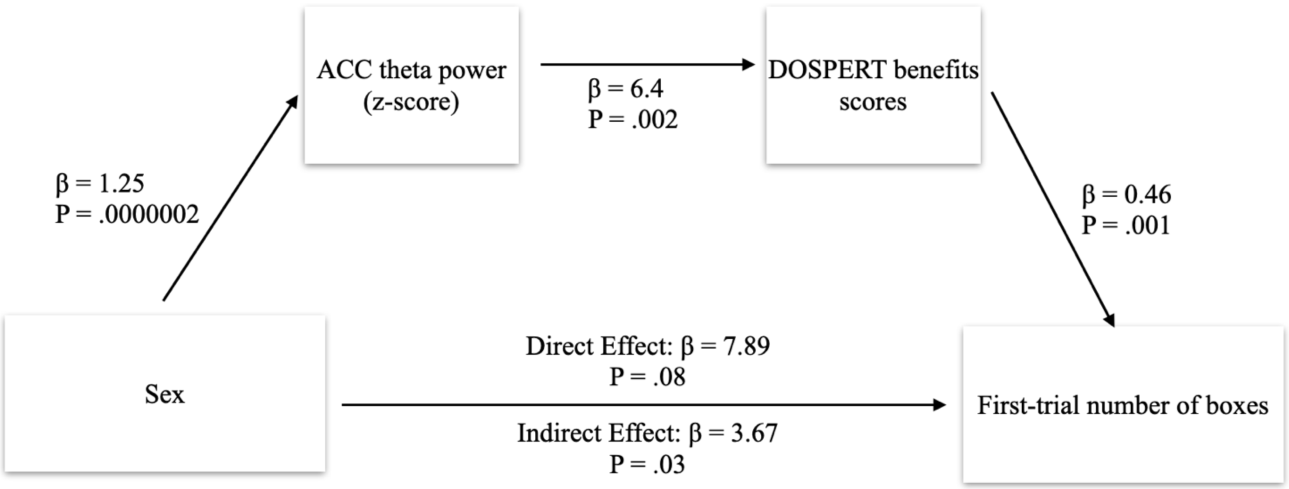


**Supplementary Figure 7.** Restricted structural equation model for sequential meditation of effects of sex on risk taking in the first trial of the game. β - regression coefficient; P - p-value of a regression coefficient. DOSPERT - Domain-Specific Risk-Taking Scale. ACC - anterior cingulate cortex.

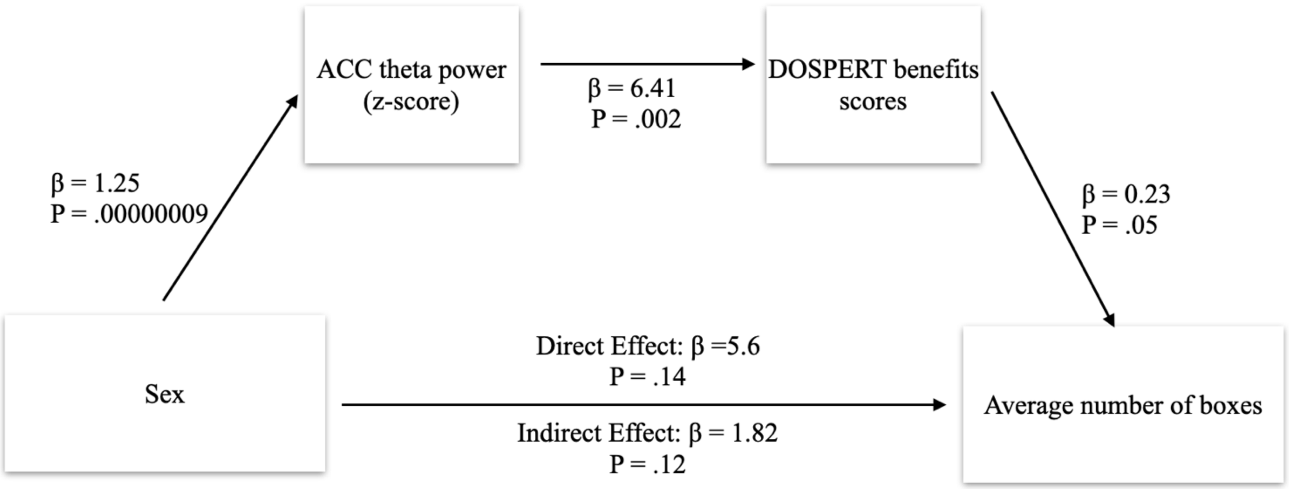


**Supplementary Figure 8.** Restricted structural equation model for sequential meditation of effects of sex on average risk taking in the game. β - regression coefficient; P - p-value of a regression coefficient. DOSPERT - Domain-Specific Risk-Taking Scale. ACC - anterior cingulate cortex.
